# Capturing the songs of mice with an improved detection and classification method for ultrasonic vocalizations (BootSnap)

**DOI:** 10.1101/2021.05.20.444981

**Authors:** Reyhaneh Abbasi, Peter Balazs, Maria Adelaide Marconi, Doris Nicolakis, Sarah M. Zala, Dustin J. Penn

## Abstract

House mice communicate through ultrasonic vocalizations (USVs), which are above the range of human hearing (>20 kHz), and several automated methods have been developed for USV detection and classification. Here we evaluate their advantages and disadvantages in a full, systematic comparison. We compared the performance of four detection methods, DeepSqueak (DSQ), MUPET, USVSEG, and the Automatic Mouse Ultrasound Detector (A-MUD). Moreover, we compared these to human-based manual detection (considered as ground truth), and evaluated the inter-observer reliability. All four methods had comparable rates of detection failure, though A-MUD outperformed the others in terms of true positive rates for recordings with low or high signal-to-noise ratios. We also did a systematic comparison of existing classification algorithms, where we found the need to develop a new method for automating the classification of USVs using supervised classification, bootstrapping on Gammatone Spectrograms, and Convolutional Neural Networks algorithms with Snapshot ensemble learning (*BootSnap*). It successfully classified calls into 12 types, including a new class of false positives used for detection refinement. *BootSnap* provides enhanced performance compared to state-of-the-art tools, it has an improved generalizability, and it is freely available for scientific use.

## 1. INTRODUCTION

The ultrasonic vocalizations (USVs) of house mice (*Mus musculus*) and rats (*Rattus norvegicus*) are becoming increasingly interesting and are investigated to better understand animal communication (for reviews see (Brudzynski, 2018; Ehret, 2018; Heckman et al., 2016)) and as a model for studying the genetic basis of autism and speech disorders in humans (Fischer et al., 2011; Scattoni et al., 2008). Rodent vocalizations are surprisingly complex and our focus here is on the USVs of house mice. Mice emit USVs in discrete units called *syllables* or *calls*, separated by gaps of silence, which have been classified into several different types by visually inspecting spectrograms (Brudzynski, 2018; Ehret, 2018; Heckman et al., 2016; Hoffmann et al., 2012; Marconi et al., 2020; Musolf et al., 2015; Nicolakis et al., 2020; von Merten et al., 2014) i.e., the squared modulus of the short-time Fourier transforms (STFT) (Oppenheim et al., 1999) (Fig. 2), or, less often, by statistical clustering analyses (Burkett et al., 2015; Chabout et al., 2017; Coffey et al., 2019; Dou et al., 2018; Hastie et al., 2009; Van Segbroeck et al., 2017). USVs are classified according to their shape and other spectro-temporal features, including the length of each syllable, their frequency content, and degree of complexity (frequency-jumps or harmonics). Our understanding of USVs has greatly improved in recent years; however, spectrograms are still usually analyzed manually (visual inspection), which is extremely time-consuming and better methods are needed for detecting and classifying USVs. Manually detecting each vocalization in many recordings can take an enormous amount of time, and though semi-automatic methods are useful, they are still time-consuming (e.g., semi-automatic detection using Avisoft SASLab Pro and manual checks requires 1–1.5 hours merely to detect 150-300 USVs (M. Binder et al., 2020), and some datasets contain tens of thousands of USVs (Marconi et al., 2020)). The time required to classify USVs takes even longer than detection, and classification is a necessary step to evaluate qualitative differences in vocalizations and to conduct analyses of USV sequences (syntax) (e.g., von Merten et al. (2014)).

Several software tools have recently become available for automating USV detection, including MUPET (Van Segbroeck et al., 2017), MSA (Chabout et al., 2017), DeepSqueak (DSQ) (Coffey et al., 2019), USVSEG (Tachibana et al., 2020), Automatic Ultrasound Detector (A-MUD) (Zala et al., 2017a), Ultravox (Noldus; Wageningen, NL) (commercial), and SONOTRACK (commercial). These tools enhance the efficiency of processing USV data, but they can generate erroneous results for several reasons. Failing to detect actual USVs (false-negative rate or FNR) can result in missing actual differences in the vocalizations of mice, and erroneous detections (false positive rate or FPR) can lead to failure to detect actual differences and generate false differences. The challenge for any USV detection algorithm is maximizing the true positive rate (TPR) while minimizing the FNR and FPR. Moreover, automatic methods can have systematic biases depending on how they are developed. For example, automated methods for detection or classification developed using only one mouse strain, one sex, one particular state, or recorded in only one context can increase both types of error (See Table 1 for the mice and recording conditions used for developing different USV detection tools if applied in other settings). Thus, automated methods can greatly enhance the efficiency of processing USV data, but it is critical that they have low and unbiased error rates. Results should be treated with caution until the error rates in the detection and classification method are evaluated.

**Table 1:**
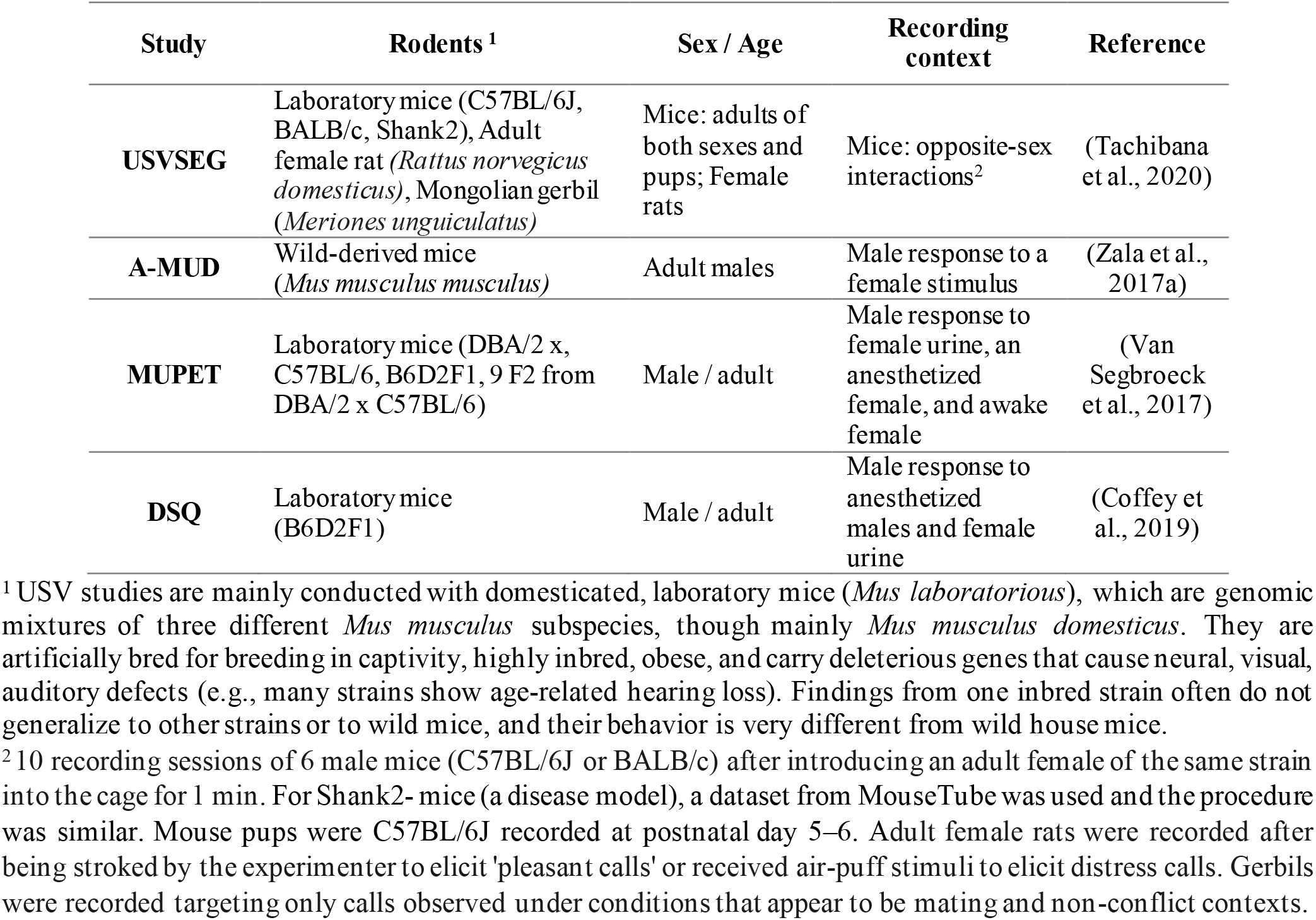
Types of rodents and recording contexts used in different studies.

| Study | Rodents <sup>1</sup> | Sex / Age | Recording context | Reference |
| --- | --- | --- | --- | --- |
| USVSEG | Laboratory mice (C57BL/6J, BALB/c, Shank2), Adult female rat (*Rattus norvegicus domesticus*), Mongolian gerbil (*Meriones unguiculatus*) | Mice: adults of both sexes and pups; Female rats | Mice: opposite-sex interactions<sup>2</sup> | (Tachibana et al., 2020) |
| A-MUD | Wild-derived mice (*Mus musculus musculus*) | Adult males | Male response to a female stimulus | (Zala et al., 2017a) |
| MUPET | Laboratory mice (DBA/2 x, C57BL/6, B6D2F1, 9 F2 from DBA/2 x C57BL/6) | Male / adult | Male response to female urine, an anesthetized female, and awake female | (Van Segbroeck et al., 2017) |
| DSQ | Laboratory mice (B6D2F1) | Male / adult | Male response to anesthetized males and female urine | (Coffey et al., 2019) |

<sup>1</sup> USV studies are mainly conducted with domesticated, laboratory mice (*Mus laboratorious*), which are genomic mixtures of three different *Mus musculus* subspecies, though mainly *Mus musculus domesticus*. They are artificially bred for breeding in captivity, highly inbred, obese, and carry deleterious genes that cause neural, visual, auditory defects (e.g., many strains show age-related hearing loss). Findings from one inbred strain often do not generalize to other strains or to wild mice, and their behavior is very different from wild house mice.
<sup>2</sup> 10 recording sessions of 6 male mice (C57BL/6J or BALB/c) after introducing an adult female of the same strain into the cage for 1 min. For Shank2- mice (a disease model), a dataset from MouseTube was used and the procedure was similar. Mouse pups were C57BL/6J recorded at postnatal day 5–6. Adult female rats were recorded after being stroked by the experimenter to elicit 'pleasant calls' or received air-puff stimuli to elicit distress calls. Gerbils were recorded targeting only calls observed under conditions that appear to be mating and non-conflict contexts.

Only five studies to our knowledge have compared the performance of USV detection algorithms: (1) M. Binder et al. (2020) compared MSA and Avisoft for detecting USVs emitted from different strains of mice (C57BL/6, Fmr1-FVB.129, NS-Pten-FVB, and 129). They concluded that Avisoft outperformed MSA for C57BL/6 and NS-Pten-FVB strains, but these two methods performed similarly for strain 129. Thus, there are strain-specific differences between these two detection tools. (2) In another study, M. S. Binder et al. (2018) compared the quantity of USVs detected by Avisoft to those detected by Ultravox (2.0) and reported significant differences in USV detection and weaker than expected overall correlations between the systems under congruent detection parameters. (3) Van Segbroeck et al. (2017) compared MUPET and MSA for detecting USVs emitted by B6D2F1 males from MouseTube (“MouseTube,”) and found that these methods generated similar call counts and spectro-temporal measures of individual syllables. (4) Coffey et al. (2019) compared MUPET, Ultravox, and DSQ for detecting USVs by analyzing the TPR and precision (the ratio of detected true USVs to false positives). For this purpose, they manipulated a recording from MouseTube in two ways to gradually degrade its quality. In the first experiment, increasing levels of Gaussian white noise were added to recordings, and DSQ outperformed MUPET and Ultravox in terms of TPR and precision in all Gaussian noise levels. In the second experiment, real noise was added to recordings, and DSQ again outperformed MUPET in terms of precision and Ultravox in terms of precision and TPR. (5) (Zala et al., 2017a) compared the performance of Avisoft and A-MUD (version 1.0) in identifying USVs of wild-derived *Mus musculus musculus*. They concluded that the latter method is superior in terms of TPR and FPR. Zala et al. (2020) have since provided an updated version of A-MUD, which overcomes previous difficulties in identifying faint and short USVs.

Our first aim here is to systematically compare the performance of four commonly used USV detection tools, MUPET, DSQ, A-MUD, and USVSEG, and we addressed three main questions:

1. How does the performance of these detection methods compare to each other? Previous studies indicate that A-MUD outperforms Avisoft, which outperforms MSA; MSA is comparable to MUPET and DSQ outperforms MUPET and Ultravox. To our knowledge, no study has systematically compared the performance of A-MUD and DSQ, nor evaluated more than two of these methods together, except for (Coffey et al., 2019), which compared DSQ, MUPET, and Ultravox.
2. How does the performance of these detection methods compare to the ground truth (i.e., detection by trained researchers)? Evaluation of detection methods rarely include a positive control (e.g., manual detection), though this is necessary to obtain absolute versus relative estimates of performance (e.g., see (Zala et al., 2017a)). For example, M. Binder et al. (2020), M. S. Binder et al. (2018), and Van Segbroeck et al. (2017) compared Avisoft and MSA, Ultravox and Avisoft, and MUPET and MSA only based on the number of USVs detected by each of the two methods, no comparisons were made with the ground truth. Coffey et al. (2019) used about 100 manually detected USVs as ground truth for comparing DSQ, MUPET, and Ultravox.
3. How well do USV detection tools generalize and perform when using data that differs from the training set (by generalization or out-of-sample error)? To our knowledge, only one study (M. Binder et al., 2020) has tested whether USV detection methods generalize to other strains (i.e., Avisoft and MSA), and only one study has compared MSA and MUPET for different recording conditions (males vocalizing in response to female urine, an anesthetized female, and awake female) (Van Segbroeck et al., 2017). Van Segbroeck et al. (2017) and Coffey et al. (2019) only used the recordings from B6D2F1 and (Zala et al., 2017a) from wild-derived *Mus musculus*. Consequently, it is unclear how well current detection methods perform whenever applied to new recordings that differ from the data used to develop and evaluate the tool. This problem is well known in the machine learning community and there are particular approaches towards this “transfer learning” (Pan et al., 2009). Thus, addressing these three questions is central to evaluating the performance of USV detection methods.

To compare the performance of these USV detection tools, we used recordings of house mice, including both domesticated laboratory mice (*Mus laboratorius*) and wild-derived house mice (*Mus musculus musculus*), and we used recordings made under different social contexts and recording conditions. To evaluate the absolute performance of these models, we applied a new dataset of manually detected USVs as ground truth with a total of 3955 USVs. The FPR is problematic for existing tools when analyzing recordings with unwanted disturbing sounds (false positives (FPs)), i.e., non-USV sounds generated because of poor recording instruments, movements of the mouse (and bedding), and social interactions during recording. Low-SNR recordings usually occur when mice are recorded with bedding in their cage and especially during social interactions, as this provides a much more natural environment for the animals. False negatives are, of course, problematic as those represent data that are just purely lost for the subsequent analysis. Signal detection theory predicts that there is an inevitable trade-off between FP and FN in the detection step (Wiley, 1983). Using a refinement step, we can set the parameters of detection such that it errs on the negative rather than the positive set, as FPs can be deleted in the refinement step. To remove FPs, MUPET and DSQ, therefore, include a preliminary detection refinement step using either an unsupervised approach, which groups data based on similarity measures rather than manually labeled USVs (both approaches), or a supervised approach, which requires manually labeled USVs for training a classifier (DSQ and (Smith et al., 2017)). Our preliminary evaluation found that DSQ outperforms MUPET in the detection refinement step (using the K-means clustering (Kanungo et al., 2002)), however, its performance differs depending on the different data. Thus, we designed a method better suited to deal with the problems mentioned above and we, therefore, compared the ability of DSQ and our classifier to detect FPs, as this is a critical step for accurate USV classification.

Classification poses an even greater challenge than detection. First pilot approaches for a similar evaluation of classification tools made it clear to us that there is potential for improvement here. Therefore, we developed an enhanced method for automatic classification, of USV syllable types. This can be achieved through unsupervised (Chabout et al., 2017; Coffey et al., 2019; Dou et al., 2018; Hastie et al., 2009; Van Segbroeck et al., 2017) and supervised (Coffey et al., 2019) classifiers. The advantage of unsupervised classification (‘clustering’) is that it does not require a predefined number of classes or manually labeled observations. The number of classes is based on the information contained in the dataset rather than the researchers’ assessment. However, these clusters do not always match those classified by researchers and it is unclear how they are perceived by mice (see Conclusions). In contrast, supervised classification (‘classification’) methods require labeled data in which USVs are classified by researchers for training a classifier (machine learning), and they have higher accuracy compared to clustering approaches (Goudbeek et al., 2008; Guerra et al., 2011). To our knowledge, only a few studies have used supervised methods for classifying mouse USVs: (1) Vogel et al. (2019) classified USVs from C57BL/6J mice into 9 classes, including ‘s’, ‘ui’, ‘c’, ‘f’, ‘up’, ‘d’, ‘c2’, ‘c3’, and ‘c’, using Random Forest (Breiman, 2001), an ensemble learning classifier of decision trees. To provide input, 104 features had first been extracted for 25-high signal-to-noise-ratio (SNR) instances from each class, and their classifier yielded a classification accuracy of 85%. (2) (Coffey et al., 2019) developed a classifier (in DSQ) based on Convolutional Neural Networks (CNNs) (Krizhevsky et al., 2012), which was trained on 56000 USVs acquired from B6D2F1 mice (MouseTube dataset). Using interpolated spectrogram images, it categorizes USVs into 5 default classes: ‘split’, ‘ui’, ‘rise’, ‘c’, and ‘c2’. (3) We (Abbasi et al., 2019) classified the elements detected from adult wild-derived house mice (*Mus musculus musculus*) into the classes ‘c2’, ‘c3’, USVs without jumps (‘no-jump’), and FP. In this work, the supervised CNNs was trained using 1200 samples and fed by 2D Gammatone filtered spectrograms (GSs), adapted to the frequency range of mice. The evaluation of its performance showed a macro-F1 score of 90±2.7%. (4) Recently, (Premoli et al., 2021) classified USVs of mice into 10 classes using different machine learning methods. The classes included ‘c’, ‘h’ (i.e., ‘c’ with additional calls of different frequencies), ‘c2’, ‘up’, ‘d’, ‘ui’, ‘s’, ‘f’, ‘c3’, and ‘composite’ (i.e., two harmonically independent components). They used 48,669 USVs of NF-kB p50 knock-out mice (B6; 129P2-Nfkb 1tm 1 Bal/J) and control wild-type mice (B6; 129PF2). Avisoft was used for USV detection. They compared the performance of CNNs fed by spectrogram images and different classical machine learning algorithms (including support vector machines) fed by 20 features. The features were obtained by Avisoft. They concluded that there is a ‘significant’ advantage using images, which contain the entire time/frequency information of the spectrogram (78.8% accuracy), rather than a subset of numerical features for classifying USVs (73.9% accuracy).

Since the generalizability of USV classifiers has never been investigated (unlike methods for classifying bird vocalizations (Brandes, 2008)), it is not known how well the current methods can classify USVs for novel datasets. So again, for this task, a systematic evaluation on a new dataset neither used for training nor for the testing is interesting. We identified three key factors that can reduce the performance and generalizability of USV classifiers:

1. Noise is a potential problem for classification, as for detection, but this issue has not received sufficient consideration. Some methods only used high-SNR data for developing their models and to improve their classification performance (e.g., (Vogel et al., 2019), (Coffey et al., 2019), and (Premoli et al., 2021)). This step results in reduced performance for newly recorded low-SNR recordings (Wu et al., 2008), which are common in practice, as argued above. This problem is exacerbated if the model is developed using predefined features extracted from spectrograms (e.g., see (Vogel et al., 2019)), as the extraction of these features from low-SNR signals already introduces high variance.
2. Imprecise USV detection generates follow-up classification errors. As the main output after detection is usually the time and frequency range of USVs, the classification will only include the region of the spectrogram limited to the detected minimum and maximum USV frequency (Coffey et al., 2019; Vogel et al., 2019). Our investigations, however, revealed that faint portions of USVs are often not included inside this window, leading to significant errors in feature estimation and classification.
3. Limited training and evaluation inflate model performance. The performance of any model is over-optimistic whenever the same type of data (same mouse strain or recording contexts) is used for the model development and also its evaluation (Abbasi et al., 2019; Premoli et al., 2021; Vogel et al., 2019). Using such a limited training set conceals the model’s shortcomings in dealing with different strains or recording conditions, but surprisingly, no previous studies have considered this issue.

Thus, to develop new and improved methods for USV classification, we aimed at the following principles:

1. Develop the first classifier based on the CNNs algorithm, which is accurate even with noisy (low-SNR) data.
2. Use the full time-frequency images based on the entire frequency range and reduce the dimensionality (and thereby the computational load) using Gammatone filters applied to the spectrograms.
3. Compare our new method with DeepSqueak (DSQ), which is currently the state-of-the-art classification tool, and evaluate it using USVs recorded under different conditions and from different mice strains than the conditions and strains used in the training step.

## 2. DATA and METHOD

### 2.1. USV data

#### 2.1.1. Subjects

The data used in this study was first divided into two meta-sets: we have used one development set (DEV) to develop, train and test the developed detection and classification method. To test the generalizability of the methods we use an additional evaluation (EV) set. For a direct test, as well as estimating the meta-parameters of the classifier, using stratified 8-fold cross-validation, the DEV dataset was further divided into three subsets including DEV_train, DEV_validation, and DEV_test (see Table 1). We report the performance of the proposed classifier in Sections 3.2 and 3.3 over the DEV_test dataset. The DEV dataset (Zala et al., 2020; Zala et al., 2017a) combined two pre-existing datasets: the first dataset was from 11 wild-derived male and 3 female mice (*Mus musculus musculus*) recorded for 10 min in the presence of an unfamiliar female stimulus (Zala et al., 2017b). In the second data set, 30 wild-derived male mice (*M. musculus musculus*) were recorded for 10 min in the presence of an unfamiliar female on 2 consecutive days, first sexually unprimed and then sexually primed (Zala et al, unpublished data). These were F1 and F2 descendants from wild-caught mice, respectively, which for brevity, we refer to as ‘wild mice.’

The EV dataset consists of two datasets, and a part was obtained from wild mice (‘EV_wild’) (as in DEV), but under different conditions (Marconi et al., 2020). The vocalizations were obtained from 22 sexually experienced adult wild-derived (F3) male *M. musculus musculus* (Marconi et al., 2020). Male vocalizations were recorded without and also during the presentation of a female urine stimulus over three recording weeks, one time per week and each time for 15 minutes. To evaluate classifier performance, we used three arbitrarily chosen recordings out of these 66 recordings, and manually classified them for this study. The other part of the EV data is taken from the MouseTube dataset used for developing DSQ (‘EV_lab’) (B6D2F1 mice recorded by Chabout et al. (2015)) and two arbitrarily selected recordings were sampled out of these 168 recordings. Although we only used a few recordings to evaluate the methods, these recordings contained a large number of USVs (Table 1). See Section Supplementary materials for more detailed information on all datasets.

#### 2.1.2. Detection

For USV detection, we applied A-MUD (version 3.2) using its published default parameters for both the DEV and the EV datasets. Because FPs and syllables are detected during the detection process, we call the detected USVs ‘elements’ rather than ‘syllables’. The parameters that affect A-MUD performance are o1_on, o1_off and if oo is enabled, oo_on and oo_off, which are amplitude thresholds in decibel. For this study, we use two A-MUD outputs: the elements time slot and the estimated track of the instantaneous frequency over time (frequency track; FT), called ‘segment info’ (Fig. 1). We also compared A-MUD to the three other detection tools, MUPET, DSQ, and USVSEG. To ensure a comparison, where AMUD is certainly not privileged, the parameters of AMUD were fixed while those of the other approaches were optimized, through trial-and-error, i.e., we used the best parameters, which provide the highest true positive rates for each detection tool, and not the default settings. The parameters used for evaluating the different tools are presented in Table 1 in Supplementary materials.

**Figure 1.**
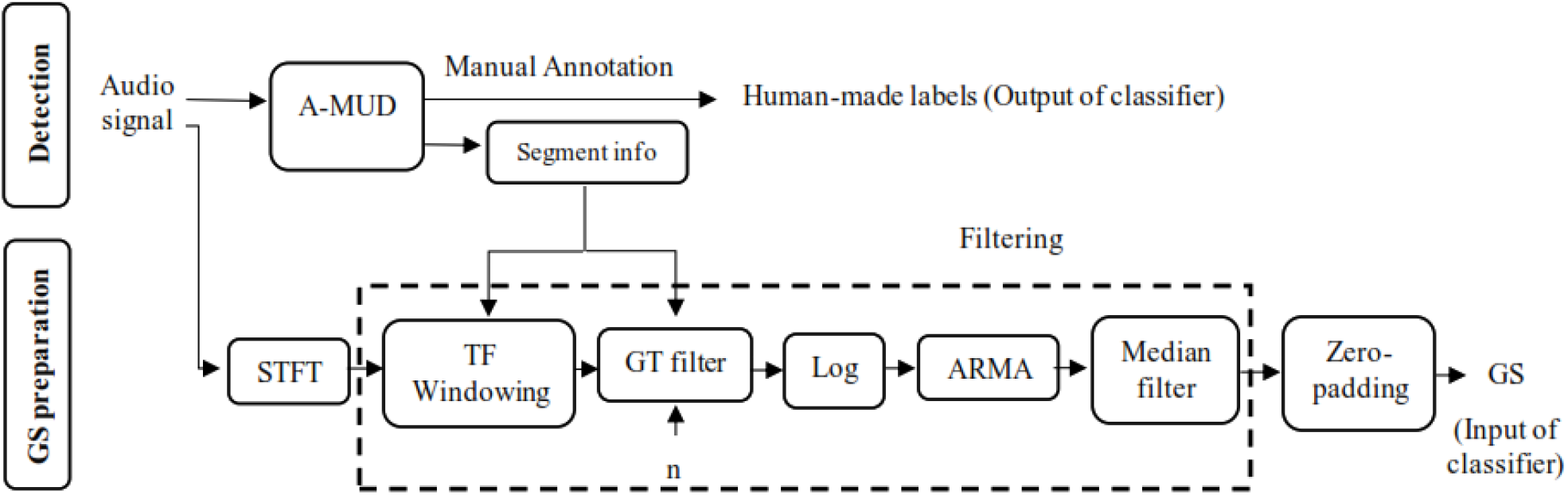
Block diagram showing the procedure for USV detection and input preparation for the classifier. ***n*** is the Gammatone (GT) filter order. STFT, A-MUD, ARMA, and GS are the abbreviation for short-time Fourier transform, automatic mouse ultrasound detector, autoregressive moving-average, and Gammatone spectrograms, respectively. TF in ‘TF windowing’ is the abbreviation for time-frequency.

Since the detection tools that we compared in this study were developed and evaluated using USVs of wild mice (A-MUD) and laboratory mice (DSQ, USVSEG, and MUPET), we also use USVs from both types of mice for our evaluation (two recordings for wild mice from the DEV and EV_wild + two recordings for the lab mice from EV_lab). The DEV_1 (1 sound file from DEV data), EV_wild_1 (sound file 1 from EV_wild data), EV_lab_1 (sound file 1 from EV_lab data), and EV_lab_2 (sound file 2 from EV_lab data) signals consist of 947, 771, 1013, and 1224 USVs, respectively.

#### 2.1.3. Manual annotation of detections

After automatically detecting all elements, the DEV dataset was manually classified into 12 classes (Figure 2), depending on the USVs’ spectro-temporal features (Hanson et al., 2012; Marconi et al., 2020; Musolf et al., 2015; Nicolakis et al., 2020; Scattoni et al., 2008; Zala et al., 2020) (Table 2 in Supplementary materials). These classes are based on frequency changes (Zala et al., 2020) (> 5 kHz increase “up”, > 5 kHz decrease “d”), on the number of components (corresponding to breaks in the frequency track; “c2” with 2 and “c3” with 3 components), on changes of frequency direction (≥ 2 changes “c”) or shape (u-shape, “u”, u-inverted shape, “ui”), on frequency modulation (< 5kHz, “f”), on time (5-10 ms, “s”, < 5ms, “us”), and harmonic elements, “h”. It is worth noting that there are 2 more USV classes, USVs with 4 “c4” and 5 “c5” components. Due to their infrequency, however, they are excluded from the training task (DEV dataset), but they are used for the evaluation step (EV dataset).

**Figure 2.**
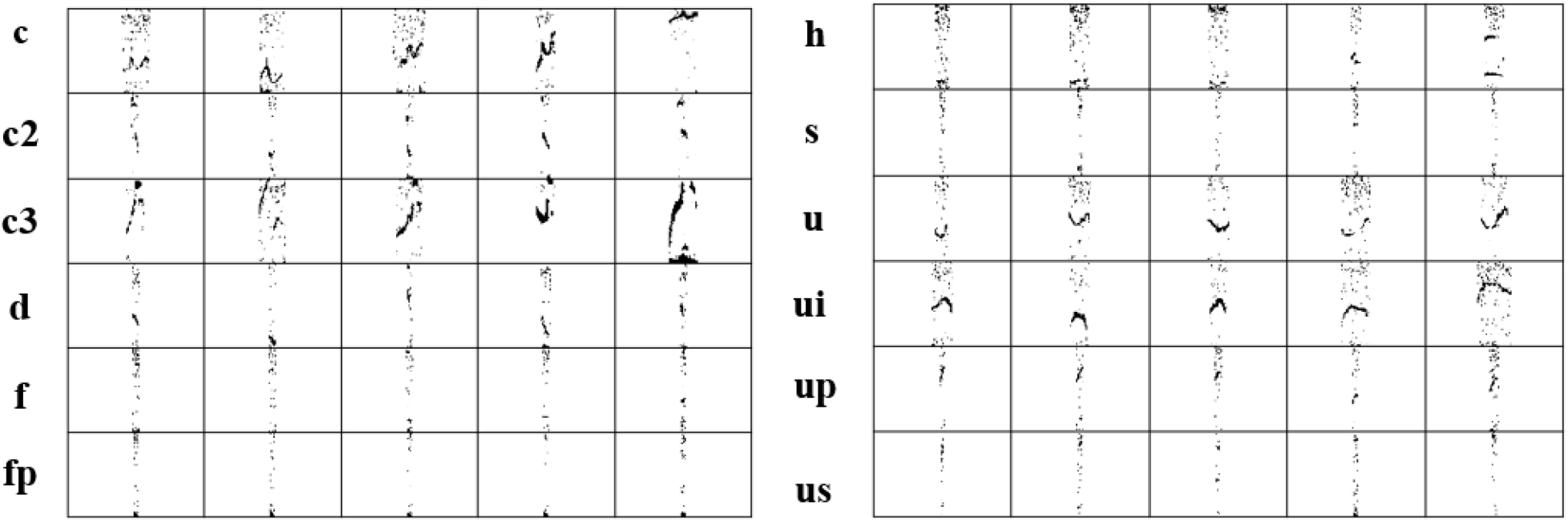
Gammatone Spectrograms (GSs) of five of five members of 12 studied classes that have the minimum Manhattan distance to other members of 12 USV classes in the development (DEV) dataset.

**Table 2.** Number of instances for each class in the different datasets.

| Data set | Number of members in each class |  |  |  |  |  |  |  |  |  |  |  |  |  |
| --- | --- | --- | --- | --- | --- | --- | --- | --- | --- | --- | --- | --- | --- | --- |
|  | **C** | **c2** | **c3** | **c4** | **c5** | **h** | **d** | **up** | **u** | **f** | **us** | **S** | **ui** | **FP** |
| DEV_train | 308 | 241 | 69 | 0 | 0 | 124 | 299 | 4343 | 298 | 1277 | 74 | 291 | 543 | 4849 |
| DEV_validation | 53 | 42 | 12 | 0 | 0 | 21 | 52 | 753 | 52 | 221 | 13 | 51 | 94 | 840 |
| DEV_test | 50 | 39 | 11 | 0 | 0 | 20 | 48 | 695 | 48 | 205 | 12 | 47 | 87 | 776 |
| EV_wild | **C** | **c2** | **split** |  |  |  | **Rise** |  |  |  |  |  | **ui** | **FP** |
|  | 20 | 224 | 334 |  |  |  | 1025 |  |  |  |  |  | 110 | 234 |
| EV_lab | 61 | 404 | 739 |  |  |  | 819 |  |  |  |  |  | 200 | 389 |

When using low-SNR recordings, or recordings with faint or short USVs, certain background noises are sometimes mistakenly detected as USVs. These errors are false positives (FPs), whereas USVs that are missed are false negatives (FNs). As mentioned above, minimizing one of these types of errors increases the other one, due to inevitable tradeoffs in signal detection (Macmillan et al., 2004). FPs are preferable over FNs, as they can be excluded in a follow-up step, and thus we included ‘FP’ as a target class. The DEV dataset contained 16958 elements including 6465 FPs in total (Table 1).

When comparing our model with DSQ, the EV data (EV_lab and EV_wild) were manually labeled into 6 classes: ‘c2’, ‘split’ (pool of ‘c3’, ‘c4’, ‘c5’, and ‘h’), ‘c’, ‘ui’, ‘FP’, and ‘rise’ (pool of ‘up’, ‘d’, ‘f’, ‘s’, ‘us’, and ‘u’). We created the classes ‘split’ and ‘rise’ because DSQ reported them together with ‘c2’, ‘c’, ‘ui’, and ‘FP’ as the output classes. The EV dataset consisted of 4500 elements including FP, of which 1947 and 2615 instances belonged to wild mice and lab mice, respectively.

#### 2.1.4. Input images for the classifier

Handcrafted, pre-determined features (such as slope, modulation frequency, number of jumps, etc.) are affected by noise, so the development of a classifier based on these features increases the error of the classification, as discussed in the Introduction. Therefore, we developed an imaged-based supervised classification built on the STFT of detected elements, followed by a set of filters and a zero-padding method (Figure 1).

After applying the time segmentation obtained from A-MUD, a 750-point Short Time Fourier Transform (STFT) (Oppenheim et al., 1999) (NFFT = 750) with a 0.8-overlapped Hamming window is applied to the signals, as shown in Figure 1. The desired information in the frequency interval of 20 kHz to 120 kHz is extracted (“TF windowing”, Figure 1). Then, following Van Segbroeck et al. (2017), a Gammatone (GT) filter bank (De Boer et al., 1978) is used to reduce the size of the STFT array along the frequency axis from 251 × 401 to 64 × 401 while simultaneously maintaining the key spectro-temporal features. This reduction can be interpreted as a pooling operator using a re-weighting step, similar to filterbanks adopted to human auditory perception (Balazs et al., 2017). Note that we adapted the frequency distribution to make our method applicable to the auditory range of mice (Van Segbroeck et al., 2017).

GT filter bank computations are provided in a MATLAB script by (Slaney, 1998). These computations were converted into the Python language for the present study. For each filter, a central frequency and bandwidth are required. The bandwidth and center frequency equations obtained in MUPET are also employed here (see Supplementary materials). In MUPET, the midpoint frequency parameter (Equation 2 in Supplementary materials) used to calculate the central frequencies was chosen as 75 kHz. The midpoint frequency can be interpreted as the frequency region where most information is processed (Van Segbroeck et al., 2017). Because the authors acknowledged that this value may not apply to all mice, we estimated the optimum value by calculating the median frequency (i.e., 63.5 kHz) from the FTs of all detected syllables, omitting FPs. Then, in a pilot test, we updated this value to 68 kHz to minimize the information loss from USVs. The central frequency was calculated based only on the DEV data. A more detailed explanation of how to determine these two parameters is given in the Supplementary materials (the Gammatone filterbank section). To further eliminate the background noise from the images, following MUPET, we calculated the maximum value between the Gammatone-filtered STFT pixels and the floor noise (10^−3^). The logarithm of the output was smoothed using an autoregression moving-average (ARMA) filter (C.-P. Chen et al., 2002) with order 1 (see Supplementary materials). Finally, a median filter (T. Huang et al., 1979) was applied to remove stationary noise. The product of the pre-processing is a smoothed, denoised spectrogram with a reduced size of 64*401, called Gammatone spectrograms (GSs). Figure 2 shows the GSs of five samples of each 12 studied classes. These samples have the minimum Manhattan distance to other members of each class.

### 2.2. CNN classifier

For our study, we used convolutional neural networks (CNNs), a particular form of the deep neural network (Goodfellow et al., 2016) first introduced by (Fukushima, 1980) and further developed by (LeCun et al., 1998). The following is a brief description of how this model works and how we implemented it.

#### 2.2.1. Classifier architecture

We used several layers: an input layer, convolution layers, pooling layers, two fully connected (FC) layers, and the output layer. The extraction of information in the CNNs is based on the 2D convolution of kernels and their receptive fields (areas on the input image determined by height and width of the kernel). The 2D convolution is performed by sliding the kernel over the entire image. The resulting matrix is called a feature map (*z_i,j_*):

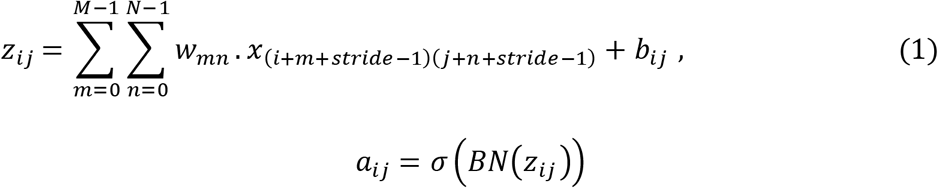

Here, *w* is the convolution kernel matrix, *b* is the bias, *x* is the input image, and *M* and *N* are the lengths and the width of the kernel. In Equation 2, the stride parameter specifying the number of pixels to shift the convolution filter is 2 for the first layer and 1 for all other convolutional layers. The batch size represents the number of training samples used for training before updating the network weights during one epoch. We trained our network with a batch size of 32 with 200 epochs. The batch-normalization layer (BN) (Ioffe et al., 2015) is calculated by normalizing the input of the layer by subtracting the batch mean and dividing it by the batch variance. The nonlinear activation function (σ) is applied to each layer output. In the current study, ELU (Clevert et al., 2015) is used for all layers except for the last one (it is softmax for the last one). After applying the activation function on the feature maps, the size of its output is reduced using a pooling layer. We used maximum pooling, which applies no smoothing and retains the key features of the image (Scherer et al., 2010). Then, the output of the last convolution layer is assigned to the FC layers to allow interactions also on a global level. The activation function of the last layer is the softmax function. The final output is calculated by taking the maximum of the softmax function output. Other activation functions (like ELU) provide an output of real-valued scores that are not conveniently scaled to be used as classifier output. However, the softmax function partitions the probability among the classes helping with the interpretation of the output, without loss of information.

The architecture of our network is shown in Figure 3. In this depiction, e.g., Conv2D (32, 3*18) denotes a 2-dimensional convolution layer with a kernel size of 3*18 and 32 filters. The FC (128) is a fully connected layer with 128 neurons. After two FC layers, a dropout layer with the probability of 0.5 is used. This step reduces the risk of overfitting (Srivastava et al., 2014). Our model has 110k parameters to be determined. The implementation is based on the Keras library (“Keras,”) (version 2.2.4) and we run the models training on the Acoustic Research Institute’s clusters with 64 GB RAM, 12-core CPUs, and NVIDIA Titan Xp GPUs, and the other with 64 GB RAM, 8-core CPUs, and NVIDIA GeForce GT GPUs.

**Figure 3.**
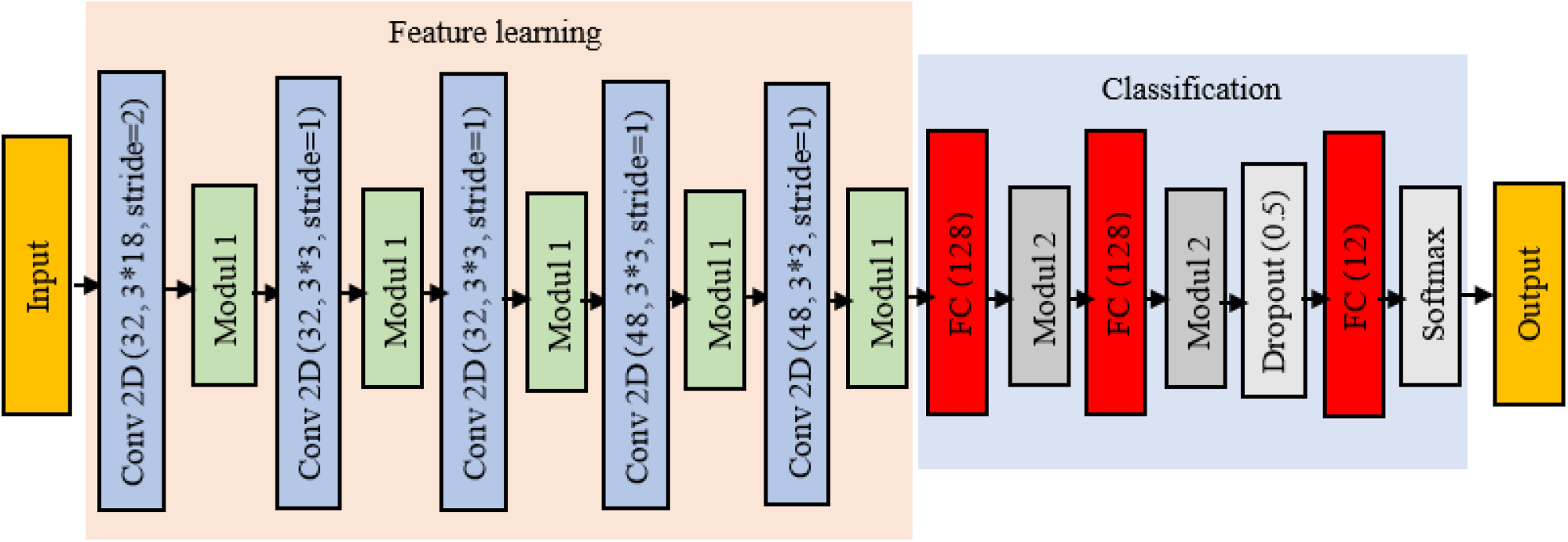
Classifier architecture. Module 1 consists of the following layers: Batch normalization + ELU + Maxpooling 2*2. Module 2 consists of the following layers: Batch normalization + ELU. Conv2D (32, 3*18) is a 2-dimensional convolution layer with a kernel size of 3*18 and the number of filters is 32. FC (128) is a fully connected layer with 128 neurons.

Data processing and analysis were conducted using Python 3.6, employing NumPy 1.16.2. Also, Sklearn 0.22.1 was used as the framework for model building and training. Figures were produced with Matplotlib 3.1.3.

#### 2.2.2. Methods for optimization and loss function

In machine learning algorithms, the general aim is to find the optimal weight to minimize the loss function. In this study, we used the categorical cross-entropy (CCE) (Goodfellow et al., 2016; Murphy, 2012), which computes the dissimilarity between the distribution of the classifier output and the manual labels. For the reduction of the overfitting (Y. Chen et al., 2016), *L*^2^ regularization (Hoerl et al., 1970), also known as Tychonov or Ridge, is added to CCE as follows,

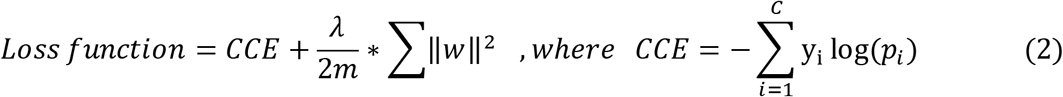

Here, *w* is the weights matrix of the CNN, ‖ · ‖ is the *L*^2^ norm, the regularization parameter λ is set to 10^−4^ and *m* is the batch size. The ground truth is denoted by y_i_ while c_i_ denotes the predicted probability of a training sample (i.e., the output of the last layer). c is the number of classes. To optimize the loss function, we used the stochastic gradient descent with Nesterov momentum (Nesterov, 1983) and we initialized the weights of the convolution and FC layers using the He-initialization (He et al., 2015).

To reduce overfitting and to promote the generalizability of the model (C. Chen et al., 2020), we performed the augmentation of the training dataset using random shifts of width and height by 10%. Other augmentation methods such as zooming and normalizing were excluded from this setup as in pilot tests, they increased the validation error of the classifier.

#### 2.2.3. Imbalanced data distribution

As shown in Table 1, the DEV_train dataset is significantly unbalanced, with 69 occurrences of the c3 and 4849 of the FP class, a typical situation in real applications of machine learning. To investigate how this uneven distribution affects the performance of the classifier, we fit the model with the original DEV_train data and it was resampled by three different approaches.

1. In the first approach, the original input data are bootstrapped *10* times to increase the generalizability and reliability of the classifier (Anguita et al., 2000; Yan et al., 2015). In each bootstrap iteration, samples are drawn from the original dataset with repetition, so some samples may appear more than once or some not at all. Then, we fitted a model for each bootstrapped dataset. The final model performance was evaluated by the average over the *10* models. Bootstrapping reduced the ratio of data imbalance from 76 to 4.
2. In the second scenario, all classes, except the classes ‘c3’ and ‘us’, which only have a maximum data number of 69 and 74, are randomly under-sampled to 124 samples.
3. In the last scenario, all classes, except FP and ‘up’, are over- and under-sampled to the number of samples of the majority class, i.e., 4849. We used the Synthetic Minority Oversampling Technique Edited Nearest Neighbor (SMOTEENN) (Batista et al., 2004) and the number of neighbors was selected as 3.

To tackle the imbalanced distribution, during the model training we also weighed the loss function inversely proportionally to the number of class members (King et al., 2001) for the original, bootstrapped, and under-sampled data using the following equation:

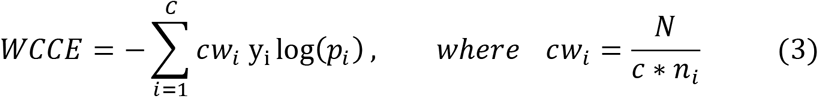

N and *n_i_* are the total number of samples and class members. CCE in equation 2 was updated to WCCE.

#### 2.2.4. Model ensemble

The weights optimized on a particular dataset are not guaranteed to be optimal (or even useful) for another dataset. At the same time, different machine-learning algorithms can lead to different results even for the same dataset. In ensemble methods (Zhou, 2012) the final output is taken from combining the outputs of different models and thus reducing the variance of the classifier output. Rather than training a model from scratch for different sets of hyperparameters, we produced 5 trained models during the training of a single model using Snapshot Ensemble with cosine annealing learning rate scheduler (G. Huang et al., 2017). They were trained consecutively, so the final weights of one model are the initial weights of the next. In this approach, the CNN weights are saved at the minimum learning rate of each cycle (Figure 2 in Supplementary materials), which occurs after every 40 epochs. To determine the best combination of these 5 models, we have cross-validated 4 approaches: 1) using the predictions of the 5th model, 2) using the average prediction from the last 3 models, 3) combining the predictions of the last 3 models by Extreme Gradient Boosting Machines (XGBMs) (T. Chen et al., 2016), and 4) combining the predictions of all 5 models using XGBMs. In explaining the third and fourth methods, instead of taking the average of the predictions (used for the second method), the predictions of the last three and five models of the DEV_validation data together with their ground truth are used for training the XGBMs. In this case, the final output of the classifier is the output of XGBMs.

Thus, to develop our classifier, these four ensemble methods were applied for each resampling approach namely under-sampling, over-sampling, and bootstrapping, and for the original data.

### 2.3. Statistical test

To determine whether the duration of USVs was statistically significant over- or under-estimated by a detection tool, a regression line (i.e., y = b0 + b1*x) was fitted between the estimated (x) and observed USV duration (y). This regression line was obtained based on ordinary least squares, which is a maximum likelihood estimator. Then, using a t-test, the P-values were calculated for the estimated intercept (b0) and slope (b1) of the regression line. These P-values assess whether the coefficients are significantly different than zero. These analyses were conducted using a Python module called statsmodels.

### 2.4. Inter-observer reliability (IOR)

Our ground truth (or ‘gold standard’) was based on manual classification, and we used two independent observers to classify USVs and to evaluate our ground truth, we evaluated inter-observer reliability (IOR). The first 100 USVs of 10 sound files were manually classified into 15 USV types by two of the authors, and both have much experience (Nicolakis et al. (2020), Marconi et al. (2020), and Zala et al. (2020)). We used five arbitrarily selected sound files from the DEV dataset and all five sound files used for the EV dataset (EV_wild and EV_lab). Both observers were blind to their respective labels and to the original labels used for the development or evaluation of *BootSnap*. The USV labels were extracted and exported into *Excel* files. The exported parameters included the start time, end time, and USV type of each vocalization. Then, the labels from both observers were aligned according to the start time of each segment. Thus, vocalizations with the same starting time were compared between the two observers. Segments that were labeled as false positive by the observers but detected by A-MUD as candidate USVs, were included and segments that were labeled as unclassified (“uc”) and were excluded from the analyses. Segments classified as the same type by both observers were scored as ‘agreement’. Segments that were either detected by only one observer or were classified into a different class were scored as ‘disagreement’. Then, we calculated the percentage of correctly classified USVs by both observers, reported as IOR. We calculated the IOR for DEV and EV data for all segments (including FPs), and when including and excluding USVs detected by only one observer and not the other (i.e., labeled as ‘missed’ USVs). In addition to the original data, we calculated the IOR and F1-score when excluding ‘s’ and ‘us’ classes, to evaluate how these two classes affected the IOR, and when pooling the original data into 12, 11, 6, 5, 3, and 2 classes, respectively, to compare the IOR and F1-score with the performance of *BootSnap* (see Table 6 and Table 7).

### 2.5. Performance statistics

The performance of the detection tools was evaluated based on TPR and FPR, which are defined as follows:

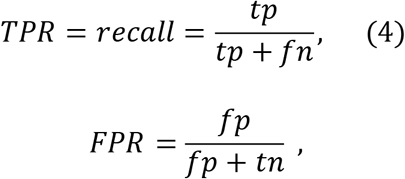

where *tp* and *fp* are true and false positives, i.e., the number of correctly and falsely detected samples of USVs, while *tn* and *fn* are true and false negatives, i.e., the correct and false number of omitted USVs.

To evaluate the performance of the classifiers, the macro F1-scores, i.e., the unweighted average of the F1-score of each class was calculated. This metric, unlike accuracy, is not affected by the imbalance distribution of the classes (Sun et al., 2009). We also used TPR and FNR (Equation 6) for producing a confusion matrix (Sammut et al., 2011).

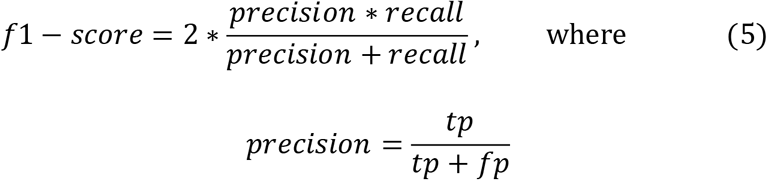

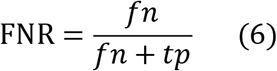

## 3. RESULTS

### 3.1. Comparing detection algorithms

Figure 4 shows the performance (TPR and FPR) of the four detection tools, MUPET, DSQ, USVSEG, and A-MUD. A-MUD was tested using its default parameters, whereas the others were implemented using the combination of parameters that provided the best results for the chosen dataset. We also compared the performance of these methods using other parameters (see Figure 2 in Supplementary materials).

**Figure 4.**
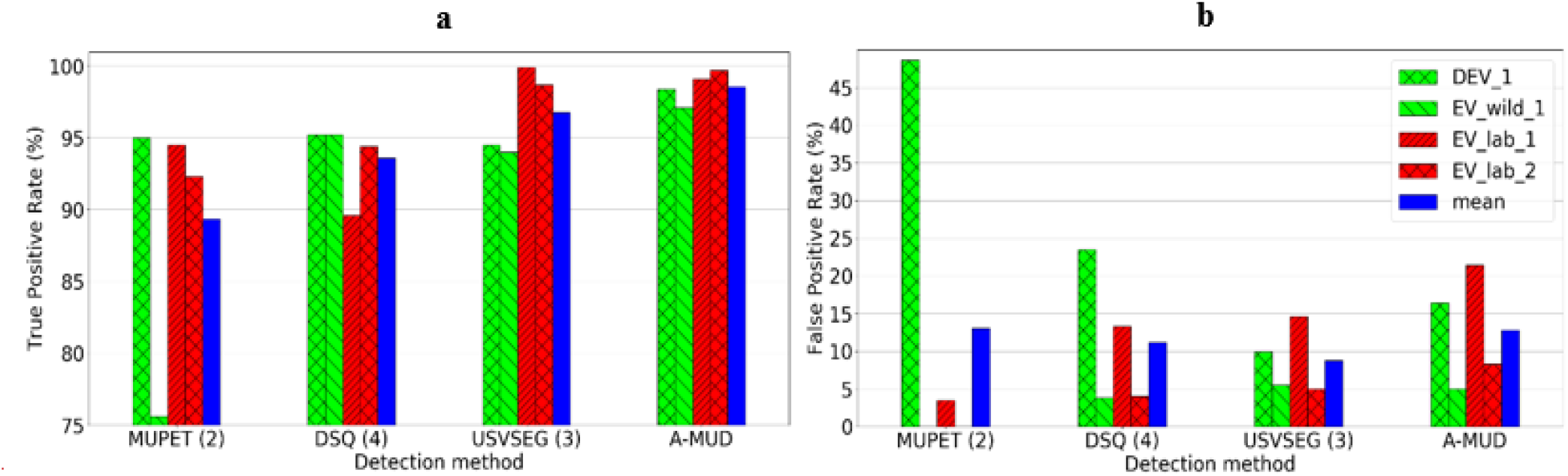
Best performance of four USV detection methods for four recordings. **(a)** The True Positive Rate shows the ratio of the number of USVs correctly detected to the total number of manually detected USVs * 100. (b) The False Positive Rate shows the ratio of the number of unwanted sounds (noise) incorrectly detected as USVs to the total number of detected elements * 100. The MUPET (2) method implemented MUPET with the noise-reduction parameter set at 5 and a minimum frequency of 30 kHz (Van Segbroeck et al., 2017). DSQ (4) used DSQ detection with the short rat call_network_v2 network with a high “recall” parameter (Coffey et al., 2019). USVSEG (3) applied USVSEG detection with the threshold parameter set at 3.5, the minimum gap between syllables at 5ms, and the minimum length of USVs at 4 ms (Tachibana et al., 2020). A-MUD was run using its default parameters (Zala et al., 2017a). The legend shows the four recordings that were compared for each method (i.e., lab mice vs wild mice for both DEV (i.e., DEV_1 and EV_wild_1) and EV datasets (i.e., EV_lab_1 and EV_lab_2) and the mean of these four recordings. DEV_1 and EV_lab_1 are examples of high-SNR recordings and EV_lab_2 is an example of low-SNR recording.

A-MUD (using the default parameters) correctly detected the largest number of USVs (TPR were all >97%), though it was closely followed by USVSEG (using the optimal parameters), and MUPET had the lowest mean TPR (<90%) (Figure 4a). A-MUD and USVSEG also provided the best performance when evaluating the detection of USVs from low-SNR recordings (DEV_1 and EV_lab_1, which include USVs from wild-derived and laboratory mice, respectively). We evaluated the performance of USVSEG using recordings of lab and wild mice and found that it has a higher TPR for lab mice. This result is likely because this method is primarily parameterized and evaluated based on recordings of lab mice. In contrast, A-MUD has a high TPR for both types of data, despite that it was parameterized and evaluated using recordings of wild mice only. The presence of faint USVs (in EV_wild_1) had little effect on the TPR for most methods, except MUPET (the TPR for this method was reduced from 95% to 75.6% when recordings contained faint USVs). When comparing FPRs, we found that USVSEG had the lowest error rates, though all four methods were similar ranging from 8% to 13% (Figure 4b). It is possible to improve the model’s performance to reduce the FPRs with an additional refinement step (see next section).

Here, we compared the estimated USV duration by USVSEG and A-MUD with the observed USV duration (i.e., manually checked and corrected USV duration). In wild mice, USVSEG underestimated the duration of USVs compared to A-MUD, which had a higher accuracy than USVSEG (Figure 5a). The duration of USVs and the mean bias values (−3.81 ms vs −0.39 ms; Figure 5c) were significantly underestimated by USVSEG (see Table 3). Also, the R-squared (R^2^) and root-mean-square error (RMSE) values, which show the correlation of the predicted and observed values and the standard deviation of the prediction error, respectively, show that A-MUD estimated the duration of USVs from wild mice with higher accuracy.

**Figure 5.**
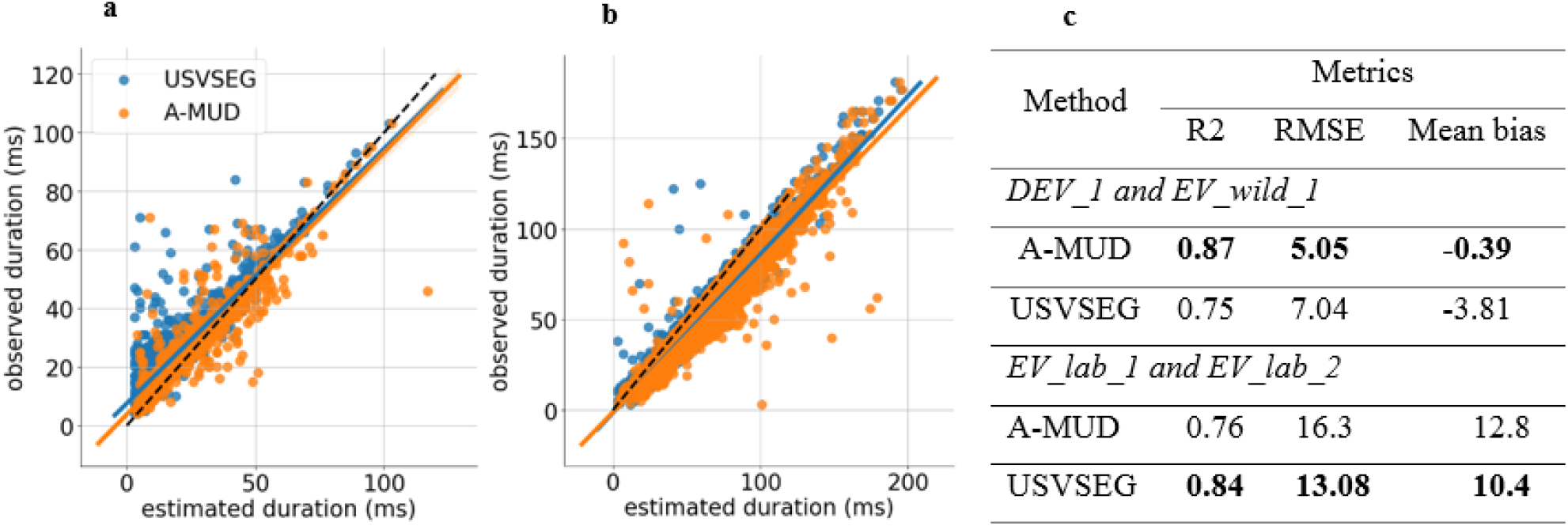
Joint plot between manually corrected (i.e., observed) and estimated duration of detected segments (by A-MUD (orange) and USVSEG (blue)) in (a) DEV_1 and EV_wild_1 data and (b) EV_lab_1 ad EV_lab_2 data. (c) Evaluation metrics for the linear regression models between observed and estimated duration of segments. The black dashed line in figures (a) and (b) is the identity line. The evaluation metrics in the table (c) are R-squared (R2), root-mean-square error (RMSE), and mean bias between observed and estimated duration of segments. Mean bias is the average difference between the estimated and observed duration of detected segments.

**Table 3.** Statistical tests comparing observed and estimated USVs duration for DEV_1 and EV_wild_1 and EV_lab_1 ad EV_lab_2 data by A-MUD and USVSEG.

| Parameters | Estimate | Std. error | t value | P value |
| --- | --- | --- | --- | --- |
| *DEV_1 and EV_wild_1* |  |  |  |  |
| Intercept_A-MUD | 3.7885 | 0.298 | 12.693 | 2.920787*10<sup>-35</sup> |
| Slope_A-MUD | 0.8941 | 0.009 | 104.872 | < 2.22*10<sup>-16</sup> |
| Intercept_USVSEG | 7.5821 | 0.318 | 23.863 | 4.396634*10<sup>-108</sup> |
| Slope_USVSEG | 0.8684 | 0.010 | 87.224 | < 2.22* 10<sup>-16</sup> |
| *EV_lab_1 ad EV_lab_2* |  |  |  |  |
| Intercept_A-MUD | -1.1628 | .045 | -2.58 | 9.8* 10<sup>-3</sup> |
| Slope_A-MUD | .084 | 0.006 | 149.926 | < 2.22* 10<sup>-16</sup> |
| Intercept_USVSEG | -1.5588 | 0.348 | -4.479 | 8*10<sup>-8</sup> |
| Slope_USVSEG | 0.875 | 0.004 | 195.535 | < 2.22* 10<sup>-16</sup> |

**Table 4.**
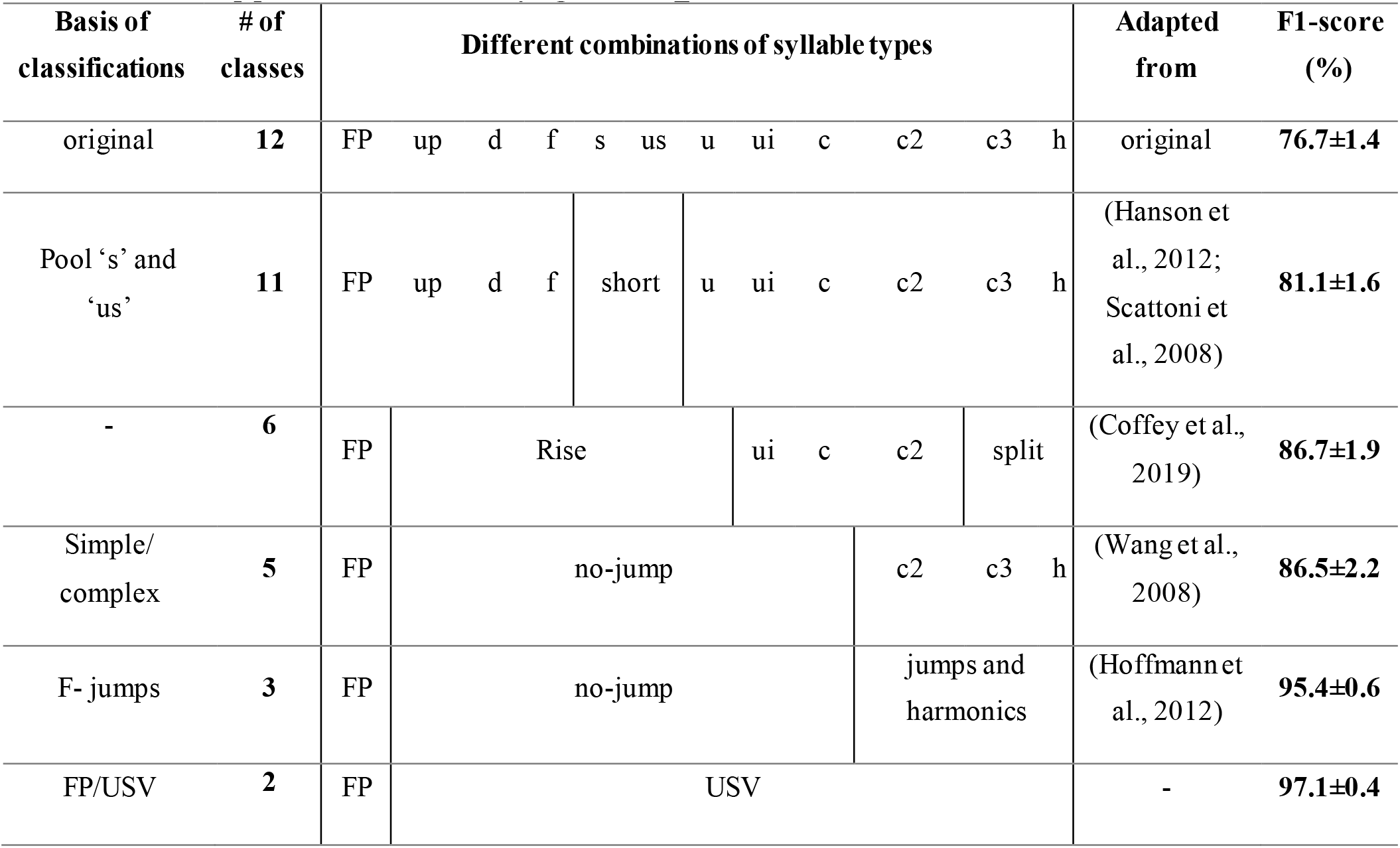
BootSnap performance in classifying the DEV_test dataset in various combinations of classes.

In contrast, the duration of USVs from laboratory mice was significantly overestimated by both methods. Here, USVSEG outperformed A-MUD, as the former had less RMSE (i.e., 13.08 vs 16.3) and higher R^2^ (i.e., 0.84 vs 0.76) than the latter. The overestimation of the duration of the USVs by both methods is probably because the USVs from lab mice were very loud and, in most cases, had a strong echo, so both methods considered these echoes as the USVs themselves. However, for the observed durations, the USVs were shortened to the end of the clear tone of the USVs.

### 3.2. Selecting the best classifier

To develop our classifier, the detected elements were first manually classified into 12 types of USVs (ground truth). In addition to the original data, three types of resampling approaches were examined (under-sampling, over-sampling, and bootstrapping) to overcome the uneven distribution between USV classes (see Section 2.2.4). For each type of resampling, four model ensemble methods were applied to the outputs, which include the predictions of the last Snapshot ensemble (‘sn’), the average prediction of the last 3 Snapshot ensemble models (‘sn_avg_3’), and a combination of the predictions of the last 3 (‘sn_xgb3’) and 5 Snapshot ensemble models (‘sn_xgb5’) by XGBMs (see Section 2.3.3). Figure 6 shows the performance of the models with different combinations of resampling and ensemble methods compared to the control run using the original data.

**Figure 6.**
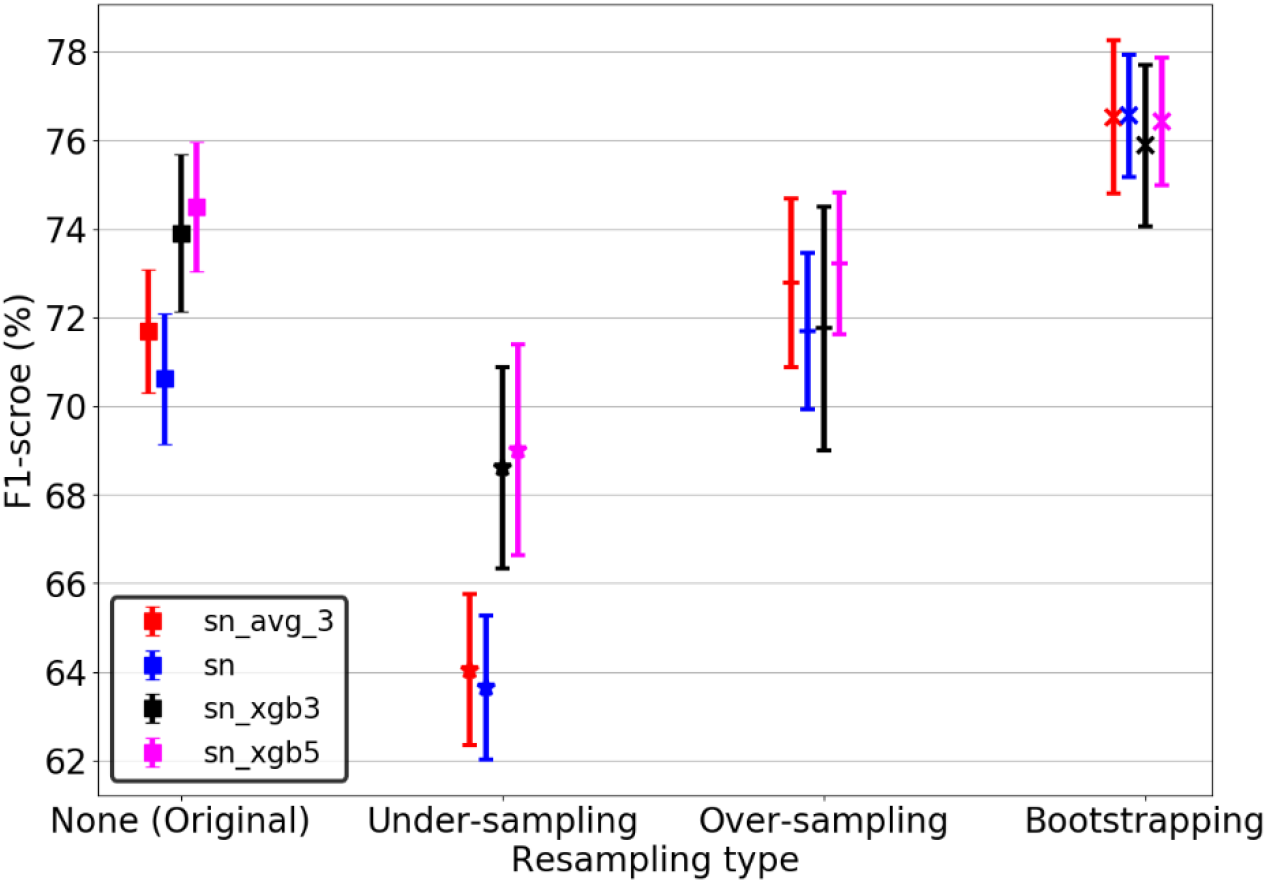
Performance of classifiers based on four resampling methods for four types of ensemble models. For each type of resampling, four ensemble models have been applied to the outputs, including the predictions of the last Snapshot ensemble (‘sn’), the average prediction of the last 3 Snapshot ensemble models (‘sn_avg_3’), and combining the predictions of the last 3 (‘sn_xgb3’) and 5 Snapshot ensemble models (‘sn_xgb5’) by XGBMs. The mean ±SD of macro F1-score of test datasets over 8-fold cross-validation are shown.

The bootstrap and under-sampling methods always had the highest and lowest average F1-score, respectively, regardless of the ensemble method. Using the last model obtained from the Snapshot ensemble gave the highest average F1-score (76.6%) with bootstrapping. ’sn_xgb5’ outperformed the other ensemble methods for the original data and two other resampling methods (under-sampling and over-sampling). The last model of the Snapshot ensemble also provided the lowest variation in bootstrapped data (1.4% STD). The differences between the ensemble methods are not large if used together with bootstrapping.

Neither the under-sampling (F1-scores = 69%) nor the over-sampling (F1-scores = 73.5%) methods, improved the performance of the model compared to the best model from the original data (F1-score = 74.5%). While this result is not surprising for the under-sampled case, the performance of the oversampling case shows that the variance is not a problem for small classes. The poor performance of the model fed by under-sampled data can be attributed to the random discard of samples and thus the deletion of useful information. The over-sampling method may have failed to improve the model performance because the images produced by the SMOTEENN are very similar to the original data (Figure 7 in Supplementary materials) leading to model overfitting. As a result, the combination of bootstrapped data and the last Snapshot model provided the best classifier (hereafter called *BootSnap*).

**Figure 7.**
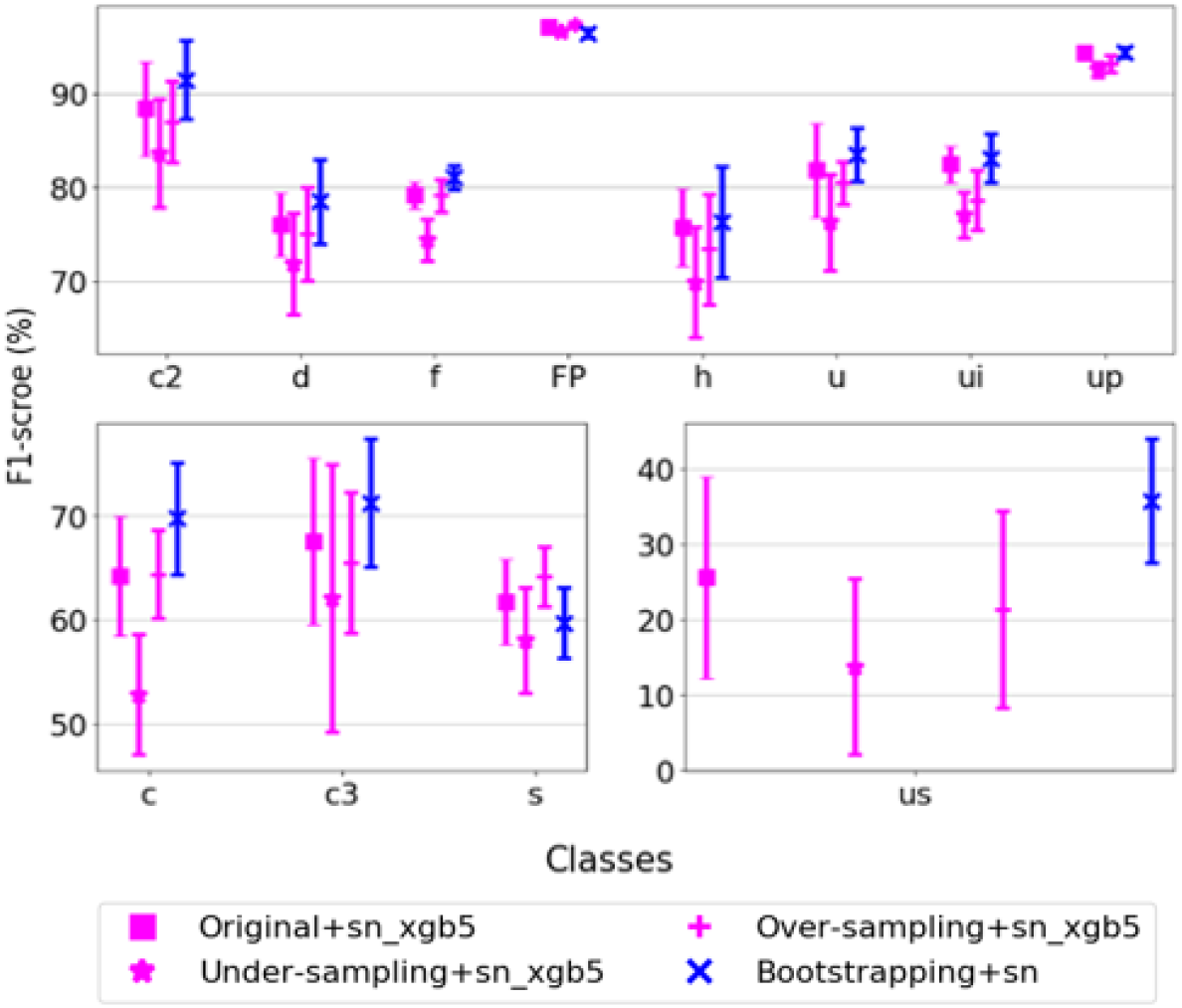
Performance of the best model for each combination of resampling and ensemble method for different USV classes. The mean ±SD of the class-wise macro F1-scores in the 8-fold cross-validation are shown.

Next, we examined the class-wise performance of the best model for each combination of resampling and ensembling method, including original + ‘sn_xgb5’, under-sampled + ‘sn_xgb5’, over-sampled + ‘sn_xgb5’, and bootstrapped + ‘sn’ (*BootSnap*). As shown in Figure 7, *BootSnap* improved the F1-scores of classes ‘c’ and ‘c3’ by about 5% and class ‘us’ by about 10%. The number of classes ‘c3’ and ‘us’ in the original data is lower than in other classes, and bootstrapping seems to effectively increase the number of class members used during the model development. For classes, ‘c2’, ‘d’, ‘f’, and ‘u’, *BootSnap* increased the average macro F1-score by about 2%-3%. The classes ‘FP’, ‘h’, ‘ui’, and ‘up’ in the original + ‘sn_xgb5’ and *BootSnap* models have approximately equal average macro F1-score. Somewhat surprisingly, the average macro F1-score of the classes ‘h’ and ‘ui’ did not increase by bootstrapping, so it seems that the number of these data points is sufficient for our method. It appears that only for the class ‘s’ bootstrapping did not help, but the abundance of class members of ‘up’ and ‘FP’ in the original data defused the effect of bootstrapping. The average macro F1-score of *BootSnap* in the class ‘s’ is about 2% less than in the model fed by the original data.

*BootSnap* also reduced the variation in the macro F1-scores for almost all USV classes, and the largest reduction in variation was for classes ‘u’, ‘c3’, and ‘us’. However, the classes ‘us’ and ‘c3’ had the highest macro F1-score STD in all resampling methods; a result that might be due to the very low number of samples in these two classes (99 and 93 members respectively).

### 3.3. Evaluating BootSnap for classifying USVs

To evaluate the performance of *BootSnap* for different types of USVs, we generated a row-wise normalized confusion matrix (or error matrix) (Sammut et al., 2011). To prepare this matrix, we used the manual annotations and predicted labels from *BootSnap* of the test dataset (of 8-fold).

This matrix shows that non-USVs (‘FP’) were classified with the highest recall (94%), which indicates that our model can successfully detect most falsely identified signals, and exclude them from further processing. It also shows that 40% to 92% of different types of USVs were accurately classified. The lowest recall was the ‘us’ class, and more than 40% of ‘us’ were mistakenly labeled as class ‘s’ and 14% of the total members were assigned to the class ‘FP’. The classification of ‘ s’ syllables (76%) was much more accurate than ‘us’, and the highest FNR value of this class (‘s’) belongs to the class ‘us’. The misclassification of these two classes can be attributed to the use of the USVs length as the only feature used for manual classification, which is not reliable (‘us’ also shows much lower inter-observer repeatability in manual classification than other classes; see Figure 6 in Supplementary materials). Class ‘c3’ had the second-lowest recall (63%), and most of its FNs were found with the classes ‘h’ (17%), ‘c2’ (9%), and ‘c’ (5%). These errors were due to the occurrence of harmonic patterns or faint jumps in the class ‘c3’. The class ‘c’ had the third-lowest recall (67%), despite having a high number of members. The problem is that ‘c’ syllables were often mis-assigned due to their similarity in the spectrograms to ‘ui’, ‘u’, and ‘up’ types, which resulted in the highest FN rates in these three classes. Examination of the misclassified members of the class ‘h’ indicates that they were often assigned to the class ‘f’. The highest portion of FNR (17%) of the class ‘c3’ is found with the class ‘h’. The FNR of the class ‘h’ is 5% with class ‘c3’. In other words, the members of the class ‘c3’ are much more likely to be mistaken as the class ‘ h’ than vice versa. It is because harmonic patterns are frequently seen with the second element (out of three elements) in the class ‘ c3 ‘, whereas the opposite rarely occurred in our recordings. The explanation might be because the ‘h’ has always only one element (+ the harmonic) and the “c3” has three elements.

As shown in Figure 2, members of the class ‘d’ resemble the members of class ‘f’, which resulted in the class ‘d’ having the most FNs with the class ‘f’. While there is no distinguished pattern of FNs distribution in other classes, it is important to note that FNs of the classes ‘c2’ and ‘c3’ mostly occur among themselves. Thus, the performance of the classifier is improved after pooling the ‘c2’ and ‘c3’ classes, as we show next.

### 3.4. Inference classification

Since it is unclear whether and how mice classify USVs, we report the performance of the best classifier (*BootSnap*) based on the different number of classes proposed in previous studies (Table 2). It is important to note that, unlike previous studies, we considered FP as a target class. Since *BootSnap* was trained using 12 classes, we pooled different types of calls in various combinations, especially for the most similar types of syllables, to compare its performance with existing literature treating other numbers of classes. This comparison provides some insights into the classification of types of USVs by researchers.

The number of USV classes studied here ranged between 2 and 12 different types. As expected, classifying all 12 classes provided the lowest F1-score (76.6±1.4%). In the next step, the classes ‘us’ and ‘s’, which differ only in their duration, were pooled to form a new class, labeled ‘short’. By combining these two classes, we found a significant increase in the F1-score (81.1 ± 1.6%). In addition, by combining these two classes, a significant number of ‘us’ and ‘s’ types, which were mistakenly assigned as each other (Figure 6), were correctly classified as ‘short’. In the next step, the classes ‘up’, ‘d’, ‘f’, ‘s’, ‘us’, and ‘u’ were pooled to form the class called ‘rise’, and the classes ‘c3’ and ‘h’ were included in the class ‘split’. Aside from the class ‘u’, a common feature between classes pooled into ‘rise’ was having no changes in their frequency direction. These classes were mostly false positives in the 12-member classification, and thus, after pooling, the F1-score significantly increased to 86.7±1.9%, compared to the 11-class classification.

Then, according to Wang et al. (2008), the number of classes was reduced to five. We pooled the classes ‘ui’, ‘c’, and ‘rise’. These classes have no jumps in their spectrograms and thus the pooled new class was labeled ‘no-jump’. Also, the classes ‘h’ and ‘c3’, which were pooled in the previous step into the class ‘split’, were separated again, but unlike the previous steps, the F1-score decreased (ca. 0.2%). This result might have been due to the separation of classes ‘h’ and ‘c3’ causing a large number of members of the latter class to be classified in the former class (Figure 5 in the Supplementary materials). In the next step, all the members of the classes ‘c2’, ‘c3’, and ‘h’ were pooled into the class ‘jumps and harmonics’ and compared with the classes ‘FP’ and ‘no-jump’. As mentioned before, all the FNs of the classes ‘c2’ and ‘c3’ were from each other (Figure 8), and as a result, pooling them in one class yielded an F1-score of about 95.4±0.6%. Finally, we classified syllables and FP into two separate classes, and this simple binary classification, which was mostly used in the USV detection step, was able to differentiate USVs from FPs with an F1-score of 97.1±0.4%. These results again show how the performance of *BootSnap* depends upon the type of USV, and that pooling certain classes results in better accuracy.

**Figure 8.**
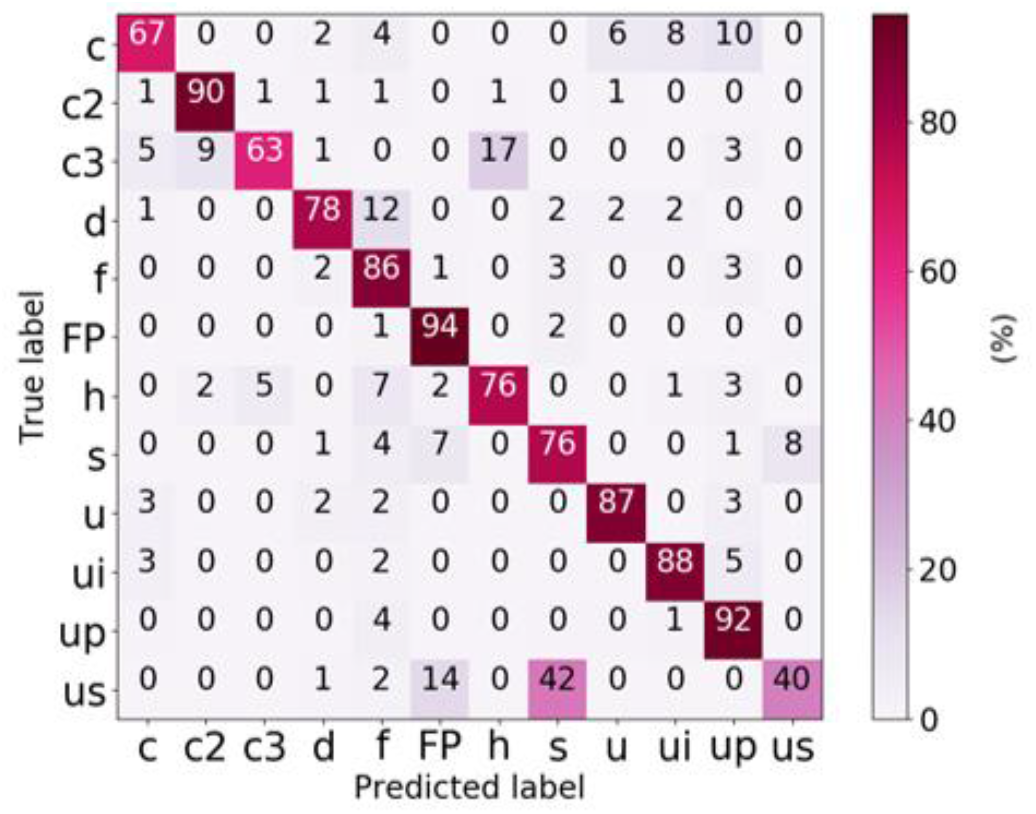
Confusion matrix of a 12-class classification using *BootSnap*. The main diagonal represents the recall of each USV class. The other values in each row are FNRs, which indicate the percentage of each class of USVs incorrectly labeled or classified.

### 3.5. Comparing *BootSnap* and DSQ: transferability to new datasets

We compared the performance of *BootSnap* to DSQ, which we consider to provide the state-of-the-art classification tool, and we used the EV_wild and EV_lab signals (Table 3). *BootSnap* predictions were pooled into 6 classes, which included ‘rise’, ‘split’, ‘ui’, ‘c2’, ‘FP’, and ‘c’ (DSQ reported them as the output classes). DSQ distinguishes FPs from USVs using a post hoc denoising network (Coffey et al., 2019) before the classification step. For comparison, we considered FP as one of DSQ’s final output. Since *BootSnap* was developed based on 8 folds, we used the mode of 8 predictions to compare it with the DSQ output. It is also important to note that A-MUD was used to detect USVs in both algorithms to provide a fair basis for comparing the classification step in DSQ and *BootSnap* (this improved the average detection rate of DSQ by 5%).

As expected, *BootSnap* and DSQ performed better for the types of mice used for training the models (wild and lab mice, respectively; Table 5). DSQ had an F1-score of 41% for wild mice and 49% for lab mice. Similarly, *BootSnap* had an F1-score of 67% and 64% for wild and lab mice, respectively. Nevertheless, *BootSnap* outperformed DSQ for both types of mice overall. In terms of class-wise performance, *BootSnap* performed better in nearly all the classes (‘c’, ‘c2’, ‘split’, ‘FP’, and ‘ui’, with higher F1-scores of 32%, 14%, 2%, 43%, and 54 % for the EV_wild and higher F1-scores of 14%, 50%, 10%, 11%, and 12 % for the EV_lab). DSQ outperformed *BootSnap* for the EV_lab for one class, ‘rise’. The reason for the superior performance of *BootSnap* in classifying ‘c2’ and ‘split’ classes in EV_lab over EV_wild is probably explained by the jumps that in EV_lab are stronger than in the wild mice data.

**Table 5.** Comparison of DSQ and *BootSnap* performances for the supervised classification of USVs in EV_wild and EV_lab recordings. The values of macro F1 (which is the average of F1-score over all classes) and class-wise F1-score (F1-score computed for each class) are presented.

| Scheme | macro<br>F1-score (%) | Class-wise F1-score (%) |  |  |  |  |  |
| --- | --- | --- | --- | --- | --- | --- | --- |
|  |  | c | c2 | split | FP | rise | ui |
| *EV_wild* |  |  |  |  |  |  |  |
| DSQ | 41 | 0 | 44 | 56 | 50 | 82 | 12 |
| *BootSnap* | **67** | **32** | **58** | **58** | **93** | **92** | **66** |
| *EV_lab* |  |  |  |  |  |  |  |
| DSQ | 49 | 24 | 43 | 74 | 66 | **69** | 16 |
| *BootSnap* | **64** | **38** | **93** | **84** | **77** | 61 | **28** |

Once again, an important point for developing and assessing the performance of a classifier is its generalizability, i.e., how well the model works when classifying data not used for the model development. In reviewing the above results, we observed that both DSQ and *BootSnap* had a relatively poor performance in the classification of the classes ‘ui’ and ‘c’. Further examinations showed that the decline in their performance in these classes was due to the significant difference in the distribution of new data with their training data. This difference is better seen in the three-dimensional t-SNE (Maaten et al., 2008) representation (using the initial dimension of 40, the perplexity of 50, and the number of iteration of 2000) shown in Figure 9. The F1-scores of ‘ui’ and ‘c’ classes were very low for both *BootSnap* and DSQ for lab and wild mice, still, *BootSnap* outperformed DSQ. In the class ‘rise’, the USVs of wild and laboratory mice have overlapped distribution, which was in contrast to the classes ‘ui’ and ‘c’ (Figure 8c). Thus, the performance of both models for this class was much better than for other classes.

**Figure 9.**
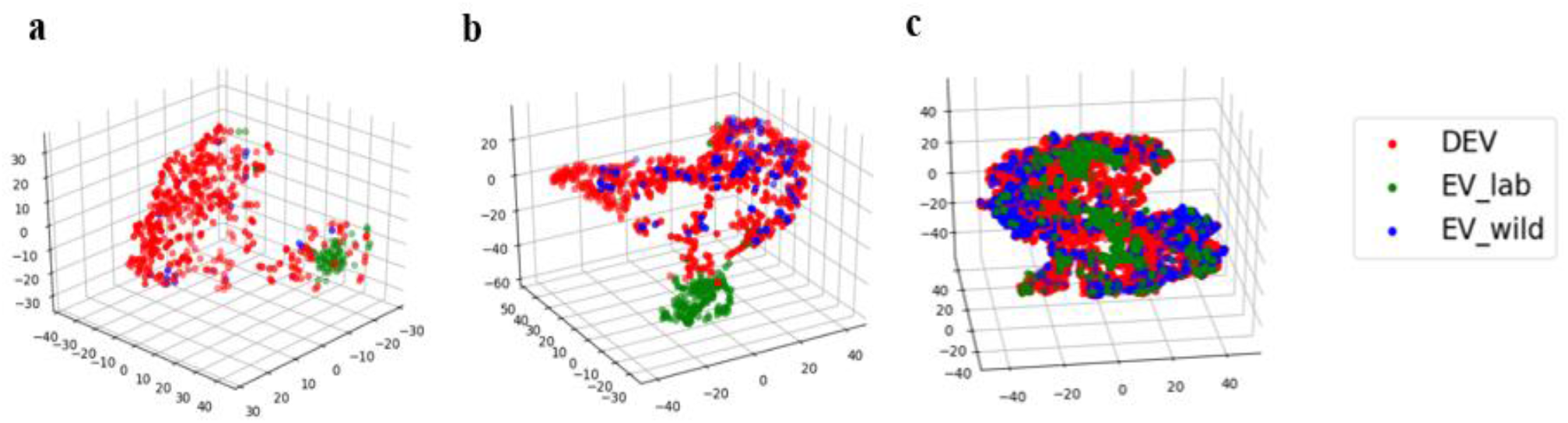
Scatterplots of USVs from three classes comparing different types of data and mice. 3-dimensional t-distributed stochastic neighbor embedding (t-SNE) representation of the classes (a) ‘c’, (b) ‘ui’, and (c) ‘rise’. Colors indicate the dataset to which USVs belong.

### 3.6. Inter-observer reliability

When calculating the inter-observer reliability (IOR), excluding ‘missed’ segments, for the DEV dataset (n = 630 segments from 5 soundfiles), we found ca. 80% agreement between two independent observers and ca. 84% agreement for the EV dataset (n = 578 segments from 5 soundfiles), when including all classes (Table 6). The removal of the ‘missed’ segments from all class combinations has a larger effect on IOR in the DEV data than the EV data. This is probably because most of the USVs in the DEV dataset have low-SNR or they are fainter compared to USVs in the EV dataset, since the EV dataset includes the EV_lab files which usually have a high-SNR (see Table 3 in Supplementary materials). So, in the EV data, the probability of error in the detection tool and observer is less due to having louder USVs.

**Table 6.** Interobserver reliability for the subsets of DEV and EV datasets. IOR values (in percentage) are given for different combinations of classes. Two IOR values are presented for each combination of classes: IOR including ‘missed’ segments / IOR excluding ‘missed’ segments.

| Interobserver reliability in various combinations of classes |  |  |  |  |  |  |  |  |
| --- | --- | --- | --- | --- | --- | --- | --- | --- |
| Dataset | Original | Excluding<br>‘s’ and ‘us’ | 12 classes | 11 classes | 6 classes | 5 classes | 3 classes | 2 classes |
| DEV | 79.5 / 85.6 | 83.6 / 87.4 | 79.5 / 85.6 | 80.6 / 86.8 | 83.8 / 90.2 | 89.2 / 96 | 89.2 / 96 | 92.4 / 99.5 |
| EV | 84 / 85.7 | 85.6 / 86.4 | 88.7 / 90.5 | 88.9 / 90.6 | 90.1 / 92 | 93.2 / 95 | 94.6 / 96.5 | 97.9 / 99.8 |

Excluding the “us” and “s” USVs increased the IOR to 84% for the DEV data (9% of the segments excluded) and to 86% for the EV data (3.6% of the segments excluded), respectively. A detailed comparison of the manual classification by the two observers (Figure 6 in Supplementary materials) showed that the USV types “us”, “s”, “up”, “u”, “h”, “c”, “c3”, “c2”, and “ui” in the DEV dataset and “us”, “s”, “up”, “h”, “c4”, “c5”, and “ui” in the EV dataset accounted for the highest disagreement between observers. The disagreement for the type “us” was likely due to detection error since “us” USVs have <5 ms duration and might not be detected by another observer in noisy recordings. If there is a disagreement in the length of USVs (due to faint USVs or background noise) between observers, an “us” might be classified as “s” and “s” USV might be classified as “d” or “us”. We observed a low number of “s” and “us” types when analyzing the EV dataset especially within the recordings from laboratory mice (9% of “us” and “s” in the DEV dataset compared to 3.6% in the EV dataset). Additionally, there can be disagreement between the USV types “up” and “ui”. This error is likely to occur due to the threshold of 5kHz to measure the frequency modulation and used to distinguish between “up” and “ui”. USVs with upward frequency modulation of >5 kHz (“up”) often ends with a slight downward frequency modulation, which can be close to 5 kHz. USVs often have a lower amplitude at the start or the end of the vocalization, and sometimes it can be difficult to measure the exact frequency modulation in a spectrogram. In summary the main misclassifications are between 1) us and s, 2) c3 and h, 3) c3, c2, and c, 4) c, ui, u, and up, and 5) d and f. Usually, the fuzzy transition between the types is the main problem in manual classification. Thus, although USV syllables are discrete, they are not all very discrete, especially when the USVs are classified into a large number of classes (e.g., more than 5 according to Table 6). These reflect that the main difficulties of *BootSnap* and manual classification are similar.

In our datasets, errors in manual classification were mainly due to (i) high background noise, (ii) duration or frequency thresholds used to define USV types, (iii) low or high amplitude of USVs (iv), and “noisy” vocalizations with many frequency jumps emitted by laboratory mice. The disagreement in manual classification of certain syllable types highlights the importance of finding a biologically relevant number of different USV classes, which can be reliably differentiated with low error rates by different observers.

Similar to the *BootSnap* F1-score, the IOR (Table 6) and F1-score (Table 7) of IOR data improved as we pooled the classes into fewer groups. For example, the IOR improved from 6 to 5 classes classification in the DEV (from 84% to 89%) and EV (from 90% to 93%) datasets. The improved IOR to 89% (DEV) and 94% (EV) after pooling all USVs with or without frequency jumps suggests that potential classification method that is more reliable between observers compared to a classification using >12 USV types. Additionally, manual classification showed an agreement of 92% (DEV) and 98% (EV) when distinguishing between USVs and false positive segments. The IOR increased to 99.5% (DEV) and 99.8% (EV) when excluding ‘missed’ segments.

**Table 7.** F1-score of the DEV_test and subsets of DEV (DEV_IOR) and EV datasets (EV_IOR) for IOR calculation. F1-score values (in percentage) are given for different combinations of classes. The numbers provided for DEV_test is the same as the numbers in Table 4. They are presented here again for easier comparison. Since we do not have ‘missed’ segments in the DEV_test data, these segments are removed when calculating the F1 score of DEV_IOR and EV_IOR datasets.

| Setting | F1-score in various combinations of classes |  |  |  |  |  |
| --- | --- | --- | --- | --- | --- | --- |
|  | 12 | 11 | 6 | 5 | 3 | 2 |
| DEV_test | 76.7±1.7 | 81.1±1.6 | 86.7±1.9 | 86.5±2.2 | 95.4±0.6 | 97.1±0.4 |
| DEV_IOR | 74.8 | 78.7 | 82.8 | 81.3 | 90 | 99.2 |
| EV_IOR | 82.8 | 83.9 | 89.7 | 84.2 | 97 | 99.6 |

Table 7 shows that in nearly all combinations of classes, F1-score of DEV_test data (calculated between ground truth and BootSnap output) is similar to the F1-score of EV_IOR and DEV_IOR datasets. F1-score of EV_IOR and DEV_IOR datasets is calculated between two observers’ labels. It can be concluded that the value of F1-score generally increases with the pooling the classes, and *BootSnap* classifies USVs with approximately equal accuracy as humans.

### 3.7. Comparing *BootSnap* and DSQ: sensitivity to new classes

One of the main performance factors of a classifier is how the classifier deals with classes for which it was not trained. The DEV data does not contain samples from two classes, ‘c4’ and ‘c5’. Therefore, to address this issue, we analyzed the performance of DSQ and *BootSnap* focusing on these two classes, which were present in EV_wild data.

The results show that *BootSnap* assigned 68% and 32% of the members of these two classes to the classes ‘c2’ and ‘c3’, respectively. It is noteworthy that both ‘c2’ and ‘c3’ classes represent jump-included USVs, which is also a prominent feature of the classes ‘c4’ and ‘c5’. DSQ assigned 3%, 13%, 46%, 3%, and 35% of the members of the classes ‘c4’ and ‘c5’ to the classes ‘c’, ‘c2’, ‘c3’, ‘rise’, and ‘ui’, respectively. Although the class ‘ui’ is relatively similar to the ‘c4’ and ‘c5’ classes based on visual inspection (see Figure 7 in Supplementary materials), the difference is that there is no jump in the class ‘ui’ to which DSQ incorrectly assigned a significant number of classes ‘c4’ and ‘c5’. Thus, we conclude that *BootSnap* uses a measure of similarity more fitted to USVs than DSQ, assigning new class samples to the most similar classes in the training data.

## 4. DISCUSSION AND CONCLUSIONS

### 4.1. Comparing USV detection tools

Our first aim was to compare the performance of four USV detection tools with each other and the ground truth (manual detection), as the detection is an important first step for classification and other analyses of USVs. Compared to previous studies, our ground truth for comparison consisted of 40 times more samples (i.e., 4000 vs 100 in DSQ), and therefore, our results should be much more robust. Moreover, we evaluated USV detection using wild mice, as well as laboratory mice, and we also compared USVs recorded on the noisy background (DEV_1 and EV_lab_1 signals) and having faint (EV_wild_1) elements. We found that A-MUD detected the largest number of actual USVs (TPRs were all >97% with its *default* parameters), and USVSEG had a similar performance (TPRs were all >94% using the adaptive optimal parameters). These two tools were better at detecting USVs from recordings with low-SNR, though faint USVs were only a problem for MUPET. USVSEG had a somewhat higher TPR for laboratory mice (99%) than wild mice (94%), and this is likely because USVSEG was primarily developed based on recordings of laboratory mice. A-MUD was parameterized using recordings of wild mice, though it still had high TPRs for both types of data, indicating that it is more generalizable than USVSEG. DSQ and MUPET had the lowest mean TPRs (94% and 89 % respectively). USVSEG had the lowest rates of false positives, though all four methods had comparable mean FPRs (i.e., between 8% – 13%). For wild mice, USVSEG underestimated more the duration of USVs compared to A-MUD (with the mean bias of −3.81 vs. −0.39, respectively). In laboratory mice, A-MUD overestimated more calls compared to USVSEG, although both methods suffer from significant overestimation of the duration of USVs.

We compared how USVSEG and A-MUD detect USVs to better understand how these methods differ. USVSEG detects USVs using the following steps:

1. it calculates spectrograms using the multitaper method, which smooths the spectrogram and reduces background noises;
2. it flattens the spectrogram using cepstral filtering, which is performed by replacing the first three cepstral coefficients to zero and subtracting the median of the spectrogram (flattening eliminates impulse and constant background noises); and
3. it estimates the level of background noise to make a threshold for the resulting spectrogram.

In contrast, A-MUD (version 3.2) detects USVs using the following steps:

1. it applies an exponential mean to the spectrograms to reduce the noise contribution;
2. it estimates the envelope of the spectrograms using 6-8 cepstral DCT coefficients;
3. it computes the segmentation parameters, which are the amplitudes (m1-m3) and frequencies (f1-f3) of the three highest peaks in the spectrum for each time position; and
4. it searches for a segment based on 4 threshold values.

The main reason for the higher performances of A-MUD (version 3.2) and USVSEG compared to MUPET is presumably because it uses flattening rather than spectral subtraction for denoising. Also, DSQ is based on training a supervised model based on a dataset (which also has high-SNR), which reduces its generalizability. On the other hand, it seems that the use of the multitaper method in USVSEG reduces the false positive rate compared to A-MUD. However, this approach in some cases leads to the disappearance of ultrashort USVs, the false detection of two USVs as a single USV, and it underestimates the duration of USVs in USVSEG. For these reasons, we utilized A-MUD for our subsequent USV detection.

### 4.2. Comparing USV classification methods

Our second aim was to develop a new method for USV detection refinement and classification and compare its performance with DSQ, and especially their relative ability to generalize to novel datasets. To develop the classifier and to overcome the uneven distribution of classes, we examined three types of resampling approaches, under-sampling, over-sampling, and bootstrapping. For each type of resampling, four model ensemble methods were applied to the outputs: the predictions of the last Snapshot ensemble; the average prediction of the last 3 Snapshot ensemble models; and a combination of the predictions of the last 3 and 5 Snapshot ensemble models by XGBMs. We found that the differences between the ensemble methods are not large if used together with bootstrapping. This result can be interpreted in such a way that the ensemble of the models derived from bootstrapped data is already compensating the uneven distribution statistically. We used bootstrapped data and the last model of snapshot ensemble as the best classifier (‘BootSnap’). The classifier had the highest errors for classifying ultrashort (‘us’) USVs mainly due to their similarity with ‘s’ USVs. These USVs do not differ qualitatively, they are not actually different syllables types, as they differ only in length. Another classification error was due to confusing ‘c’ and ‘c3’ syllables. The low recall in classifying “c3” syllable types was likely due to their small number used for training, and also because some members have a harmonic element, much like “h” types. The similarity in the spectrograms of ‘c’ to other classes as ‘ui’, ‘u’ and ‘up’ classes lead to errors in the classification of this class. On the other hand, the model classifies classes “up”, “FP”, and “c2” with a recall higher than 90% and classes “ui”, “u” and ‘f’ with a recall of more than 85%. These classes have a relatively larger number of members compared to other classes (‘us’ and ‘c3’) and their spectrograms are relatively different from each other. The overall F1-score of the model increased from 76.7% to 81.1% by pooling ‘s’ and ‘us’ classes, which resulted in a more robust classification.

We compared the performance of *BootSnap* to DSQ, which is currently the state-of-the-art classification tool. DSQ uses a 6-member syllable classification that includes ‘rise’, ‘split’, ‘ui’, ‘c2’, ‘FP’, and ‘c’ types (i.e., a simpler classification approach based on 6 classes, see Table 5). USVs from wild mice as well as laboratory mice were used to evaluate the generalizability of these two classifiers. As expected, in *BootSnap* classifier, the closer the data is to the training domain, the better the overall performances. It has 85% F1-score for 6-class classification of USVs on DEV_test data (Table 4), but about 65% F1-score for EV datasets. We found that our new classification method outperformed DSQ in nearly all aspects, including USVs of both the wild and laboratory mice (macro-F1 score of 66% vs 47%). This difference in performance is mainly because the DSQ classifier was developed using high-SNR data, compromising its performance with new low-SNR recordings. In contrast, we used low-SNR data to develop our classifier and aimed to enhance its ability to generalize. We also used the Ensemble learning method, which is based on the Snapshot Ensemble and Bootstrapped input data for training the classifier. In Ensemble learning, base models are combined to prevent the final model from either overfitting or underfitting, making the model more stable and generalizable.

*BootSnap* also showed better performance than DSQ in assigning new class samples to the most similar classes in training data. For example, our results show that *BootSnap* assigned all instances with more than 3 jumps (similar to those not found in the training data) to similar classes with less than 3 jumps. However, DSQ allocates 30% of these new samples to the class without any jumps. Our method also detects noise in new data much more accurately (F1-score of 93% vs. about 50% for EV_wild and 77% vs. 66% for EV_lab), and thus it is more useful for low-SNR data, which is a common challenge for USVs studies – especially studies aiming to record animals under social contexts. Another advantage is that DSQ is based on MATLAB, which requires the purchase of required licenses, whereas our method is based on Python and, thus, it is free of charge.

### 4.3. Inter-observer reliability (IOR)

To our knowledge, this is the first time that USV detection or classification tools have been evaluated that also examined the accuracy of the ground truth used to assess machine performance. According to the inter-observer reliability (IOR) results, the agreement between two observers in DEV and EV dataset was 76% and 88%, respectively. The mentioned values are related to the classification of segments into 12 classes, and, in addition to the A-MUD detections, segments which were missed by A-MUD but manually detected by one or both observers are included. A closer look at the results reveals that mislabeling members of the classes ‘us’ as ‘s’, ui’ as ‘up’, and ‘c’ as ‘ui’ and to a lesser degree as ‘up’, and vice versa, is very likely. The reason for the error in these classes is their sensitivity to the threshold (based on duration or modulation frequency) that are used in their definitions. On the other hand, in class “h”, due to the possibility of a faint harmonic element, incorrect labeling of these segments is very likely. Hence, part of the classification error of a classifier can be attributed to the error in the manual labeling of segments. However, the classes can be pooled to increase the amount of IOR (from 80% of 12-class classification to 84% of 6-class and to 93% of 2-class classification, see DEV dataset in Table 6), as this increased the F1-score of *BootSnap* (F1-score changed from 77% of 12-class classification to 87% of 6-class and 97% of 2-class classification, see Table 4). These results suggest that the error rate will depend upon the number of classes chosen for the classification, and that *BootSnap* can classify USVs with an accuracy similar to the results obtained from human inter-observer reliability.

While completing the final draft of our present manuscript, a new tool, called ‘Vocalmat’ (Fonseca et al., 2021), was published that detects and classifies USVs into 11 categories. The Vocalmat classifier is trained on the USVs of mouse pups (5 to 15 days old) of both sexes of several inbred strains, including C57BL6/J, NZO/HlLtJ (New Zealand Obese), 129S1/SvImJ, NOD/ShiLtJ (Non-obese Diabetic NOD), and PWK/PhJ (descendants from a single pair of *Mus musculus musculus*). It was developed using USVs in the frequency range of 45 kHz to 140 kHz. After contrast enhancement and applying several filters, the authors calculated the spectrogram (with the size of 227*227) of 12,954 detected elements. Its classifier is the AlexNet model (Krizhevsky et al., 2012), which was pre-trained on the ImageNet dataset. Like other studies, this classifier was not compared with other USV tools and the results on its generalizability were not provided. We evaluated the performance of Vocalmat on its test data and found that the average class-wise accuracy is 79%, whereas *BootSnap* yielded an average class-wise accuracy of 83% for classifying DEV_test elements into 11 classes. The differences in the performances of these tools could be due to differences in the test data used for evaluation.

### 4.4. Outlook

As with existing USV models, our classification method is supervised, and so if the user wants to retrain it, manually labeled data are required. On the other hand, despite the outperformance of *BootSnap* over DSQ, *BootSnap* still has difficulties with classifying new data of a complex (with no jump), u-inverted, and 1-jump including USVs. Considering that our best model is based on the bootstrap technique, naturally as the number of bootstrap iterations increases, so does the computation time. By default, 10 repetitions are considered for *BootSnap*. This means that *BootSnap* calculations will be 10 times slower than similar models. Because manual labeling of data is a difficult and time-consuming task, it is important to be able to apply a model trained on a single data source on other data sources as well. So, to further improve the generalizability of a classifier, in addition to implementing the bootstrap technique, we will increase the number of samples by using more mice recordings. We expect that this approach will increase the predictive power of our classifier.

Finally, it is important to note that the USVs of mice have been classified by human researchers based on visual inspection of spectrograms or statistical clustering models, and it is still unclear whether mice can discriminate most types of USVs. Mice can hear high frequencies and can distinguish frequencies that differ by only 3% (de Hoz et al., 2014), but there have only been few tests to determine whether mice discriminate different types of USVs. One study found that laboratory mice can be trained to discriminate simple versus complex USVs, and they also discriminated certain variations in shape and frequency (Neilans et al., 2014). A second study found that trained mice discriminate USVs depending upon their spectro-temporal similarity, and ‘classified’ complex calls and up-shapes, but not u-shaped calls (Screven et al., 2019). A third study found that mice fail to discriminate between synthetic sounds with different shapes, i.e. up- vs. down-shapes (Screven et al., 2016). The shapes of these synthesized sounds were very different from mouse USVs, however, and may have lacked characteristics critical for discrimination. Thus, future studies are needed to determine whether mice can discriminate the types of USVs proposed by researchers, and these should include recordings with normal variation of syllable types within and between each category (i.e., mice should be better able to discriminate between-versus within-syllable type variation). Until such studies are conducted, USVs classified by humans or statistical models would be more accurately labeled as *putative* mouse USVs.

## Author contributions statement

**RA**: Conceptualization; Methodology; Software; Validation; Formal analysis; Resources; Data curation; Writing – original draft preparation; Writing – review & editing; Visualization

**PB**: Conceptualization; Methodology; Validation; Resources; Writing – original draft preparation; Writing – review & editing; Supervision; Project administration; Funding acquisition

**MAM**: Validation; Investigation; Resources; Data curation; Writing – original draft preparation; Writing – review & editing

**DN**: Validation; Investigation; Resources; Data curation; Writing – original draft preparation; Writing – review & editing

**SMZ**: Investigation; Resources; Data curation; Writing – original draft preparation; Writing – review & editing; Supervision; Project administration; Funding acquisition

**DJP**: Conceptualization; Resources; Data curation; Writing – original draft preparation; Writing – review & editing; Supervision; Project administration; Funding acquisition

MAM and DN made equal contributions. SMZ and DJP made equal contributions.

## Acknowledgment

We would like to thank Anton Noll for making A-MUD outputs available.

## Ethics statement of animal experiments

There were no new experiments in this study.

## Availability of Data, Software, and Research Materials

The scripts are available on our GitHub page.

## Supplementary materials

### 1. Data

#### 1.1. Subjects

The subjects were adult wild-derived house mice (*Mus musculus musculus*), F1, F2 or F3 descendants of wild-caught mice trapped at the Konrad Lorenz Institute of Ethology, Vienna, Austria (48°12’38” N, 16°16’54”E). We used wild-derived rather than wild-caught mice to control for age and rearing conditions. Mice were weaned at 21d and kept in mixed-sex groups with ≤4 siblings per cage until the age of 5 weeks (35d). Henceforth, adult males were housed individually to prevent fighting, and females were housed in sister-pairs whenever possible. The mice were housed in standard cages with bedding, nesting material, a nest box, and a cardboard roll. Food and water were provided *ad libitum*. Housing facilities were kept in standard conditions (22 ± 2 °C and a 12:12 h white light: red light cycle, lights off at 15:00). All recordings were conducted after 15:00 when the mice are most active. We also used recordings of laboratory mice (strain B6D2F1/J) from MouseTube (Chabout et al. (2015)).

#### 1.2. Datasets

Our analyses were conducted using 169 sound files of 48 mice from four different datasets which were recorded in three different contexts or retrieved from MouseTube, respectively. These recordings were either used for development (DEV) or evaluation (EV) of the new method.

The development (DEV) was conducted using sound files of 44 individual mice from two different datasets and experiments. The first dataset in the present study consisted of 14 recordings of 10 min duration (each) from F1 mice (subjects: n= 11 males and 3 females; mean ± SD age: 204 ± 17 d; stimulus females: n = 11 and age: 181 ± 15 d), which had been socially primed by a short direct interaction with a female 1d before the recordings (n = 10 priming females, mean ± SD age: 184 ± 16 d) (Zala et al., 2017b); sex differences reported in (Zala et al., 2017b); results of priming effects reported in Zala et al. (2020)). The second dataset consisted of 10 min recordings of 30 wild-derived (F2) male mice (mean± SD age: 220± 25 d; n = 30 males; and 217 ± 30 d, n = 60 females) recorded twice over two consecutive days (Zala et al, unpublished data). The dataset included 150 sound files from 30 mice recorded over 2 days: 100 sound files of 1 min duration (10 sound files x 5 mice x 2 days = 100 files), due to setting adjustments, and 50 files of 10 min duration (1 sound files x 25 mice x 2 days = 50 files).

The evaluation (EV) was conducted using 5 arbitrarily selected files from the third and fourth datasets. The third dataset consisted of a subset (n=3 soundfiles of 5 min duration) from recordings of wild-derived mice during stimulation with a female odor stimulus (Marconi et al. (2020)). USVs were recorded from adult males (F3, generation, n = 2 males; mean ± SD age: 355 ± 65 d) recorded three times over three consecutive weeks (see below). The fourth dataset consisted of 2 arbitrarily selected sound files of 5 min duration from 168 recordings of laboratory mice (B6D2F1 mice), which were retrieved from MouseTube (Chabout et al. (2015)).

#### 1.3. Recording procedures and apparatus (socio-sexual contexts)

The mice for the first dataset were recorded for 10 min while presented with an unfamiliar stimulus female on the opposite side of a partition, which allowed visual and olfactory stimuli but not direct contact (see details in (Zala et al., 2017a)). A condenser ultrasound microphone (Avisoft Bioacoustics/CM16/CMPA) was positioned 10 cm above the subject’s compartment and was connected to an UltraSoundGate 116-200, Avisoft Bioacoustics, Germany.

The mice for the second dataset were recorded for 10 min while separated from a female stimulus by a perforated partition, and then the divider was removed allowing males to interact with the stimulus female and they were recorded for 10 min (as described in (Nicolakis et al., 2020)). The two mice were then separated again by the divider and recorded for an additional 5 min. An ultrasound microphone (USG Electret Ultrasound Microphone, Avisoft Bioacoustics / Knowles FG) was positioned 10 cm above the male’s compartment and connected to an A/D-converter (UltraSoundGate 416Hb, Avisoft Bioacoustics). This entire procedure was repeated and conducted on the next day with another unfamiliar stimulus female (Zala et al, unpublished data.). This procedure allowed us to monitor changes in vocalizations as courtship progressed over time, and the mice also obtained socio-sexual contact and experience through indirect and direct interactions. We recently found that mice significantly increased the amount of USVs (vocal performance) and the number of syllable types (vocal repertoire) after sexual priming (Zala et al. (2020)) and after the partition was removed and they began interacting directly (Nicolakis et al., 2020). For the second dataset of this study, we only used recordings during the first 10 min (with the divider) on both days (before and after sexual experience). All recordings for both datasets were conducted inside a recording chamber lined with acoustic foam.

The mice for the third dataset were recorded in a cage with bedding without any stimulus for 5 min (pre-stimulation phase), and then again for an additional 10 min while presented with female urine stimulus (as described in Marconi et al. (2020)). The urine was a 60 μl pool of thawed female urine (from 3 different unfamiliar females) presented on a cotton swab attached to the cage lid. Mice were recorded in a separate room with no observers or other animals present. This procedure was repeated for each male over 3 consecutive weeks, resulting in a total of 66 recordings. For USV recordings, an ultrasound microphone (USG Electret Ultrasound Microphone, Avisoft Bioacoustics / Knowles FG) was placed 10 cm above the cage and connected to an A/D converter (UltraSoundGate 416Hb, Avisoft Bioacoustics). For each male, the recording of the 10 min stimulus presentation was saved as two separate 5 min sound files to facilitate the processing of single sound files. The 3 arbitrarily selected 5 min sound files used for the third dataset in this study were all recorded during the urine stimulation.

The fourth dataset retrieved from MouseTube (Chabout et al. (2015)) originally included 10 min recordings of 12 adult male mice. Mice had 5 min control recordings during the habituation period without any stimulus inside a clean cage. Then, the males were recorded when exposed to 4 different stimuli for 5 min: fresh urine from either females or males, awake adult female, anesthetized adult female, and anesthetized adult male. Each male was exposed to the same stimulus on three consecutive days and to a different stimulus over 4 consecutive weeks (as described in Chabout et al. (2015)). For the USV recordings, ultrasound microphones (UltraSoundGate CM16/CMPA) were placed over the center of the cage in the recording box and connected to an A/D converter (UltraSoundGate 416H, Avisoft Bioacoustics). Sound files were available on MouseTube and for the fourth dataset of this study 2 sound files were arbitrarily selected from the available soundfiles. All recordings for all datasets were conducted using the RECORDER USGH software (Avisoft-RECORDER Version 4.2) with a sampling rate of 300 kHz and 16-bit format with 256 Hz FFT size for the first 3 datasets, and with a sampling rate of 250 kHz and 16-bit format with 1024 Hz FFT size for the fourth dataset, respectively.

#### 1.4. USV detection and manual classification

For all datasets, manual USV classification was conducted in STx (Balazs et al., 2000; Kasess et al., 2013). Spectrograms in STx were generated using a Hanning window with a range of 50dB, a frame of 4 ms and an overlap of 75% and the spectrogram displayed frequencies up to 150 kHz (Zala et al., 2017a), Zala et al, unpublished data, (Nicolakis et al. (2020), and (Marconi et al., 2020)). USVs and other ambiguous sounds were visually and acoustically inspected. For the first three datasets, USVs were originally labeled according to one of the 12 (first dataset) (adapted from (Musolf et al., 2015), (Hoffmann et al., 2012), (Hanson et al., 2012), as cited in (Zala et al., 2017a) or 15 (second and third dataset) USV categories (Nicolakis et al. (2020), Marconi et al. (2020), and Zala et al. (2020))) and for the fourth dataset, USVs were labeled according to 6 classes.

For the DEV datasets including the first and second experiment, the USVs were classified (or reduced) into 11 USV categories (Supplementary Table 2). The ‘uc’ and some ‘uh’ were excluded from the classification (i.e., 10.5% of the ‘uh’ from the first dataset). However, for the first dataset 89.5% of the ‘uh’ and for the second dataset all ‘uh’ were included in other USV categories if also their spectrographic shape was annotated (e.g., if a USV was originally labeled as ‘uh-up’ because it was over 91 kHz and its shape was ‘up’, it was renamed to ‘up’). The ‘c4’ and ‘c5’ were rarely detected in these sound files and therefore excluded. In summary, the DEV datasets included 11 USV classes (‘up’, ‘d’, ‘c2’, ‘c3’, ‘c’, ‘u’, ‘ui’, ‘f’, ‘s’, ‘us’, and ‘h’) and the FPs (false positives, errors due to the low-SNR recordings) to reach a total of 12 classes. The EV datasets including the third and fourth datasets consisted of 6 classes: ‘c2’, ‘split’ (pool of ‘c3’, ‘c4’, ‘c5’, and ‘h’), ‘c’, ‘ui’, ‘rise’ (pool of ‘up’, ‘d’, ‘f’, ‘s’, ‘us’, and ‘u’), and FP. We created the classes ‘split’ and ‘rise’ because DSQ (DeepSqueak) does not differentiate between individual USVs pooled in these two new classes.

**Supplementary Table 1.**
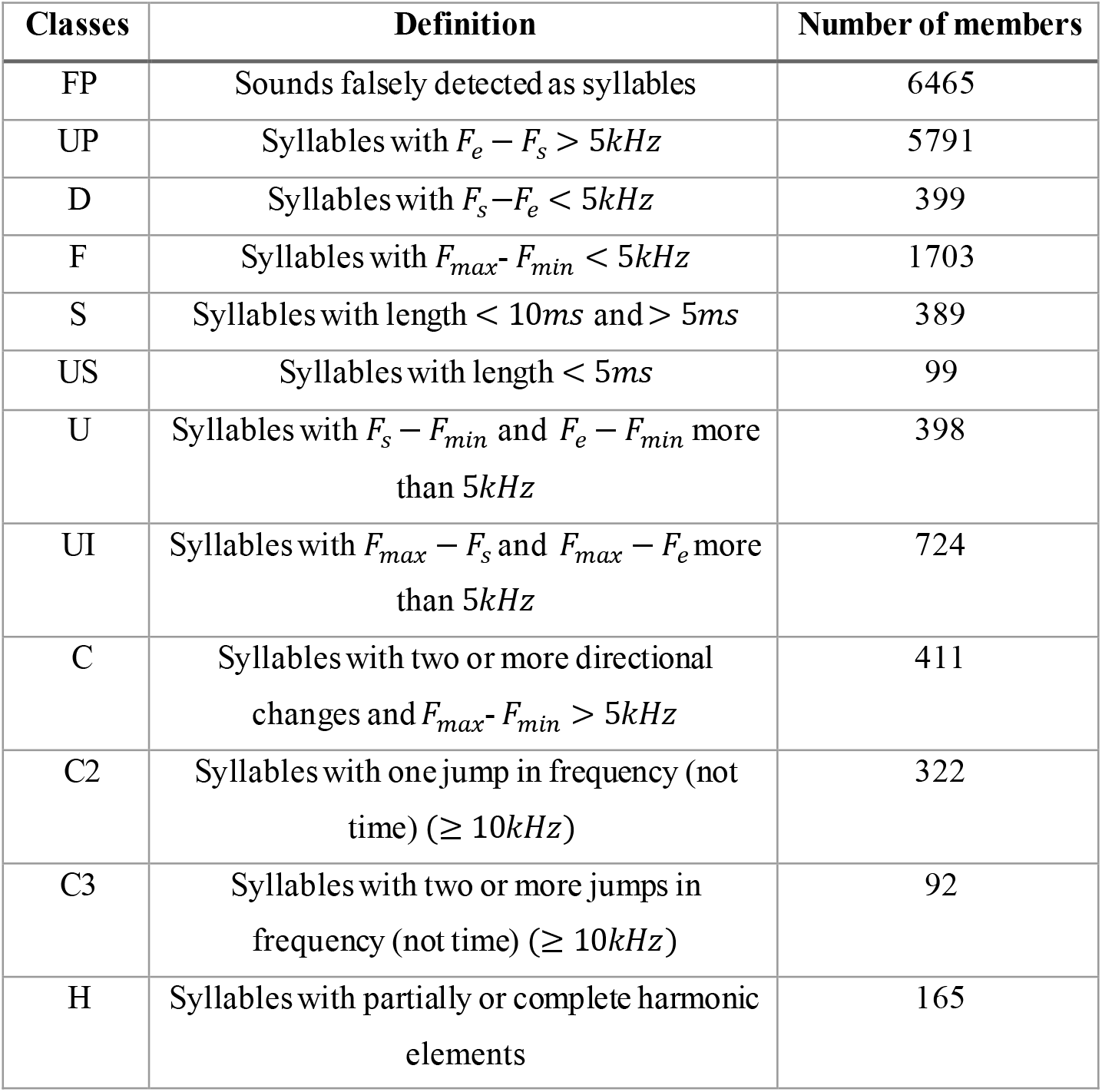
Definition of classes used in the labeling. Note that the number of members differs before and after down-sampling. ***F_e_*** is the end frequency, ***F_s_*** is the start frequency, ***F_max_*** is the maximum frequency, and ***F_min_*** is the minimum frequency. The number of members of each class corresponds to the DEV dataset.

| Classes | Definition | Number of members |
| --- | --- | --- |
| FP | Sounds falsely detected as syllables | 6465 |
| UP | Syllables with $F_e - F_s > 5kHz$ | 5791 |
| D | Syllables with $F_s - F_e < 5kHz$ | 399 |
| F | Syllables with $F_{max} - F_{min} < 5kHz$ | 1703 |
| S | Syllables with length $< 10ms$ and $> 5ms$ | 389 |
| US | Syllables with length $< 5ms$ | 99 |
| U | Syllables with $F_s - F_{min}$ and $F_e - F_{min}$ more than $5kHz$ | 398 |
| UI | Syllables with $F_{max} - F_s$ and $F_{max} - F_e$ more than $5kHz$ | 724 |
| C | Syllables with two or more directional changes and $F_{max} - F_{min} > 5kHz$ | 411 |
| C2 | Syllables with one jump in frequency (not time) ($\geq 10kHz$) | 322 |
| C3 | Syllables with two or more jumps in frequency (not time) ($\geq 10kHz$) | 92 |
| H | Syllables with partially or complete harmonic elements | 165 |

As mentioned in the main text, we compared different tools of USV detection. The following table presents the various parameters used to evaluate these tools.

**Supplementary Table 2.** The evaluated parameters for different USVs detection tools.

| Detection<br>method | Setting number |  |  |  |
| --- | --- | --- | --- | --- |
|  | 1 | 2 | 3 | 4 |
| **DSQ (Coffey et al., 2019)** | All Short<br>Calls_Network_V1 | Mouse Call_Network_V2 | Short Rat<br>Call_Network_V2 | Short Rat<br>Call_Network_V2<br>with high recall |
| **MUPET**<br>**(Van Segbroeck et al., 2017)** | noise-reduction=6<br>min-frequency=35kHz | noise-reduction=5<br>min-frequency=30kHz | × | × |
| **USVSEG**<br>**(Tachibana et al., 2020)** | threshold=4.5<br>min-length=5 ms<br>gap min =30 ms | threshold=3.5<br>min-length=4 ms<br>gap min= 30 ms | threshold=3.5<br>min-length=4 ms<br>gap min= 5 ms | × |
| **A-MUD**<br>**(Zala et al., 2017 a)** | o1-on=12 dB,<br>o1_off=10 dB,<br>min-length=5 ms |  |  |  |

### 2. Gammatone spectrograms preparation

In speech, unsupervised methods such as Non-negative matrix factorization (NMF) (Févotte et al., 2011; Lee et al., 1999) are used to reduce the size of the spectrogram while preserving the time-frequency information. Using NMF, the audio signal spectrogram is approximated using the weighted sum of the basis unit functions, so that the basis unit functions and their weights are non-negative. According to studies, the basis unit functions (or spectral bases) obtained from NMF are very similar to the human cochlea’s biological and perceptual time-frequency resolution (Fletcher, 1940), as well as perceptual scales, such as the Mel (Stevens et al., 1937) and bark scales (Zwicker, 1961). In MUPET, NMF has been applied on the USVs spectrogram to reduce their size along the frequency dimension. The NMF output is the product of spectral bases matrix, which are band-pass filters and are modeled by Gammatone band-pass function, and their weights, which are the spectral magnitude associated with the corresponding filter. To preserve most information and reduce the computational load, the number of spectral bases has been selected as 64. A regression is fitted to the peak frequencies of the base functions to determine the center frequencies and bandwidths of the gammatone filters, which are as follows:

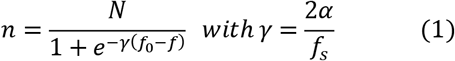

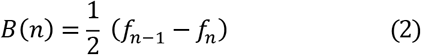

f_s_ is the sampling frequency (i.e., 300 kHz) and N corresponds to the chosen number of filters in the filterbank (i.e., 64). *f*_*n*–1_ and *f_n_* are the central frequency of n-1^th^ and n^th^ Gammatone filter, and B is Gammatone filter bandwidth.

The midpoint frequency (f_0_) and the slope variable (α) were initially obtained from the MUPET script (f_0_)=75 kHz and α=14.2). We changed these two parameters (to 68 kHz and 16, respectively), so that all 64 Gammatone filters are generated in the range of 20 kHz to 120 kHz. The variable slope was set based on trial and error as 16. f_0_ is modified based on the mean frequency of the USVs in our data at which most USVs occur. For the calculation of the mean frequency of USVs, we used the frequency track of USVs, which was explained in the Methods section (Input images for the BootSnap). The middle Gammatone filter has the lowest bandwidth (i.e., 0.57 kHz) due to the high number of USVs in midpoint frequency. Other Gammatone filters, which are symmetrically distributed, have higher bandwidth (i.e., between 0.57 kHz and 6.6 kHz) due to the smaller number of USVs in frequencies lower and higher than midpoint frequency.

In the next step, the Gammatone filters are applied as weighted summation kernel to the STFT of USVs subsequently thresholded. This threshold is 10^−3^, so the maximum value between the Gammatone-filtered STFT pixels and the floor noise (10^−3^) was calculated. The output is logarithmically transformed and, then, it is smoothed using an Auto Regression Moving-Average (ARMA) filter (Box et al., 1970) with order 1.

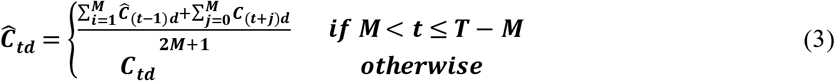

The variable 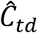 is the spectrum filtered by ARMA, the *C_td_* is the spectrum filtered by the Gammatone filterbank, and *M* is the filter order (Van Segbroeck et al., 2017). Finally, the median filter is applied to remove stationary noise from *C_td_*. Then zero padding is applied to produce images of USVs with the same size of 401*64. 401 is the width of images, which is related to the maximum duration of USVs (i.e., 200 ms) and 64 is the number of Gammatone filters. Supplementary Figure 1 shows (a) the probability distribution of USVs Frequency track values used to update f0, (b) the frequency response of 32 Gammatone filters, (for simplicity 32 filters were plotted), and (c) two examples of the USVs spectrogram before (top row) and after applying the Gammatone filter and post-processing steps discussed above (bottom row).

**Supplementary Figure 1.**
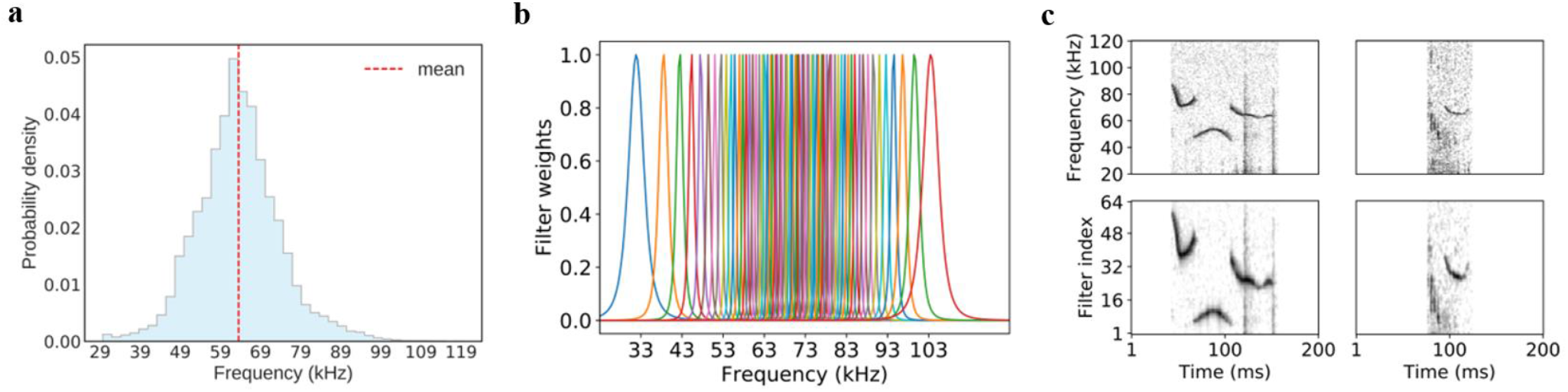
(a) Distribution of USVs Frequency Track (FT) values, extracted by A-MUD. The FT values are related to all detected syllables, omitting false positives. (b) The frequency response of 32 Gammatone filters (we have used 64 filters, but for simplicity 32 filters are plotted here) at the frequency range of 20 kHz to 120 kHz. (c) Two examples of the USVs spectrogram before (top row) and after applying the Gammatone filter and post-processing step (bottom row). This image shows that by applying the preprocessing steps on the spectrogram, although the size of the images is reduced from 251 × 401 to 64 × 401, the important information of the USVs is not lost.

### 3. Classifier

As mentioned in the original text, the learning rate used in this study is based on cousin learning rate, which is defined as follows.

**Supplementary Figure 2.**
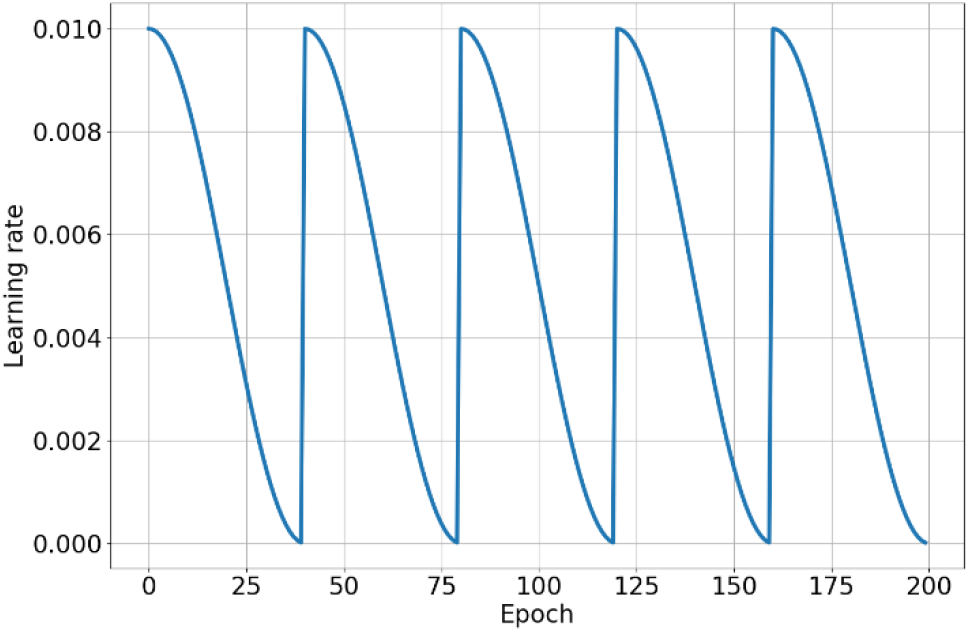
Schedule scheme used for the learning rate. Using this scheme of learning rate, the final weights of the model at every 40 epochs are the initial weights of the model in the next epoch. In this approach, the CNN weights are saved at the minimum learning rate of each cycle, i.e., at every 40 epochs.

### 4. Result

#### 4.1. Detection

In the main text, we compared the performance of 4 USV detection tools (USVSEG, A-MUD, DSQ, and MUPET) in a setting (i.e., input parameters) of which the selected parameters lead to their best performance for the four-given data (DEV_1, EV_wild_1, EV_lab_1, and EV_lab_2). Here, we compared the performance of these methods using all the combinations used for their parameters (Supplementary Table 1).

**Supplementary Figure 3.**
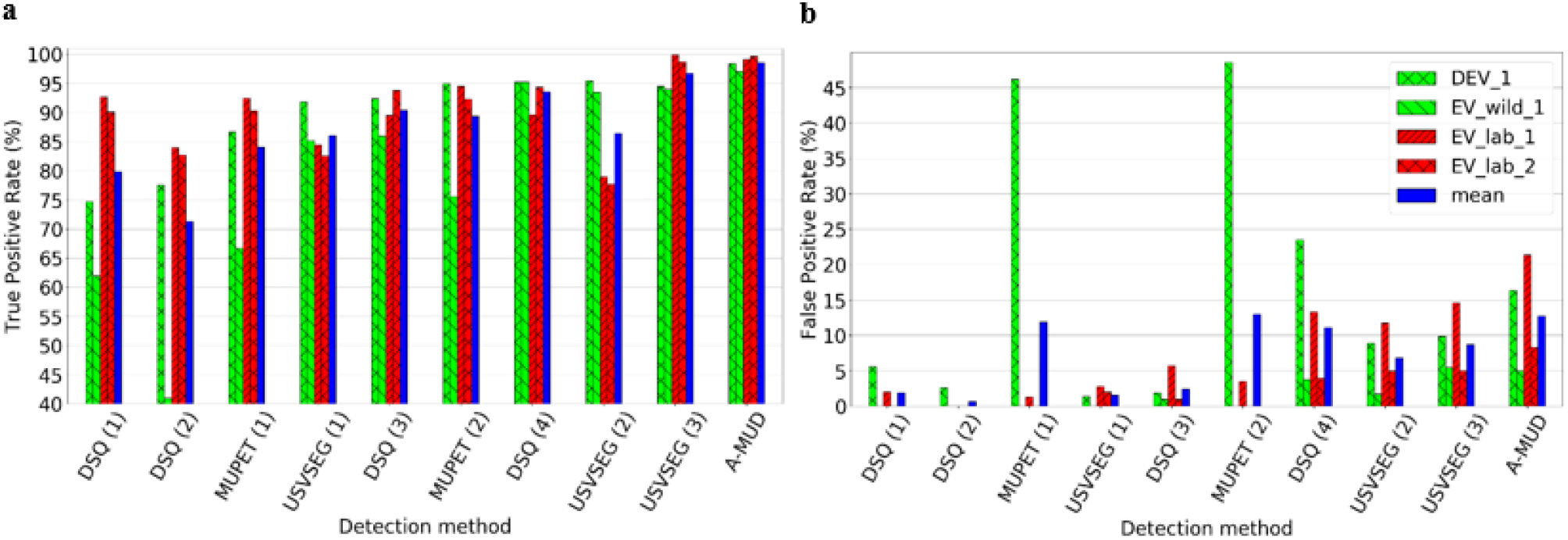
a) true positive rate (TPR) and b) false positive rate (FPR) of detection tools. If we want to compare the best performance of each detection tool with the best performance of others, A-MUD and with a slight difference, USVSEG are in the first and second places, followed by DSQ and MUPET. But if we do not consider the best performance of each tool (obtained using optimal parameters), this ranking will be different. In this case, A-MUD is the best tool, and DSQ (3) with the TPR of 90% and the FPR of 2.4% is superior to the other two methods. As a result, the type of parameter selected for each tool affects the superiority of their performance in the USV detection compared to others.

#### 4.2. Classification

In the model design section, we used various approaches to deal with the problem of the imbalanced datasets, including using original, down-sampled, bootstrapped, and over-sampled data. The following figure presents the over-sampled data by Synthetic Minority Oversampling Technique Edited Nearest Neighbor (SMOTEENN) presented.

**Supplementary Figure 4.**
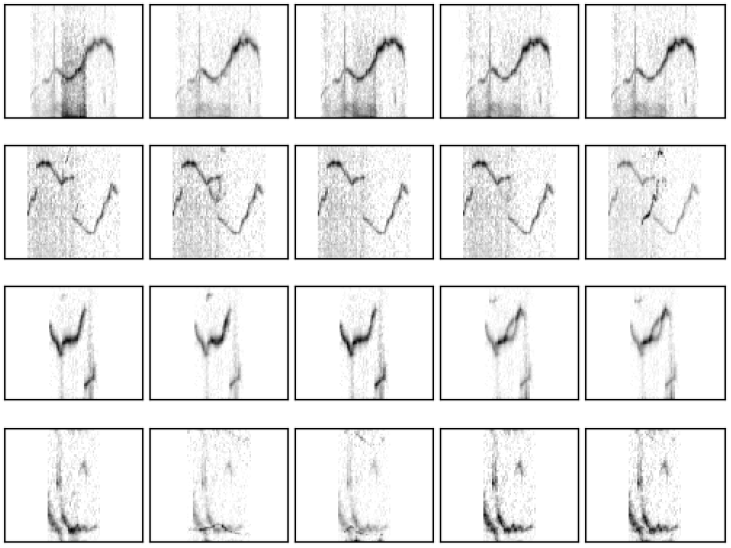
Samples produced by SMOTEENN (Batista et al., 2004). The first column from the left is the original instance and the next columns are the resampled samples. The first, second, third, and last rows are from the classes ‘c’, ‘c3’, ‘c2’, and ‘h’, respectively. The images produced by the SMOTEENN are very similar to the original data, so, compared to the original data, this method did not help to improve the classifier performance.

#### 4.3. Interobserver reliability (IOR)

In the main text of paper, we presented IOR values for various combination of classes in DEV and EV datasets. The following figures shows the normalized confusion matrix based on the labels assigned by two observers.

**Supplementary Figure 5.**
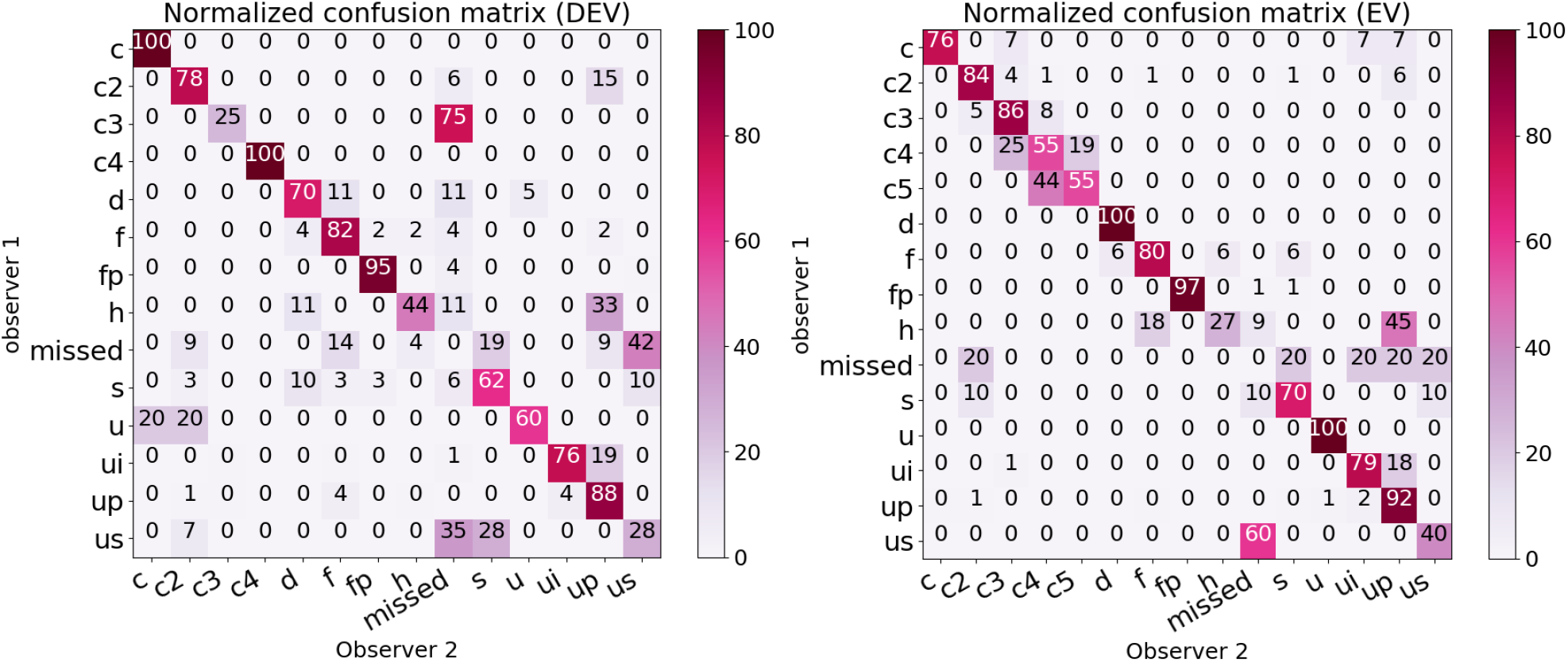
Agreement between two observers for two subsets of model development (DEV, left) and evaluation (EV, right) data. ‘missed’ segments are elements that are manually detected by only one observer. Both figures show high disagreement between the observers for both data in the ‘us’ and ‘h’ classes. In more detail, the amount of reliability in the DEV data in ‘c3’ and ‘u’ classes is very low. Differently, in the EV data, the reliability is less than other classes in ‘c4’ and ‘c5’ classes.

The table below shows the number of samples in each class in the data examined for IOR calculation.

**Supplementary Table 3.** Number of samples of each class of the observer 1 in DEV and EV subsets for IOR calculation. In DEV sub-dataset (n=5 soundfiles), there are very few samples (i.e., 2, 4, or 6) from the classes ‘c’ and ‘c4’, ‘c3’ and classes ‘u’ and ‘h’ (i.e., 6 or 10), thus the results of these classes are not very reliable. We found similar results in the EV sub-dataset (n=5 sound files) where there are very few samples from the classes ‘us’, ‘d’, ‘c’, and ‘c5’.

| Dataset | c | C2 | C3 | C4 | C5 | d | F | fp | h | missed | s | u | ui | up | us |
| --- | --- | --- | --- | --- | --- | --- | --- | --- | --- | --- | --- | --- | --- | --- | --- |
| DEV | 2 | 34 | 4 | 2 | 0 | 17 | 41 | 121 | 9 | 21 | 29 | 5 | 113 | 219 | 14 |
| EV | 13 | 64 | 79 | 36 | 9 | 8 | 15 | 75 | 11 | 5 | 10 | 14 | 53 | 181 | 5 |

#### 4.5. Comparing *BootSnap* and DSQ: sensitivity to new classes

As mentioned in the results section (Section 3.7), the performance of a model is important when dealing with a new class. Because there was no sample of the ‘c4’ and ‘c5’ classes in the DEV data, we compared the output of the BootSnap and DSQ methods when the two classes were in the EV data. The following figure shows example of members of these two classes in EV_wild data.

**Supplementary Figure 6.**
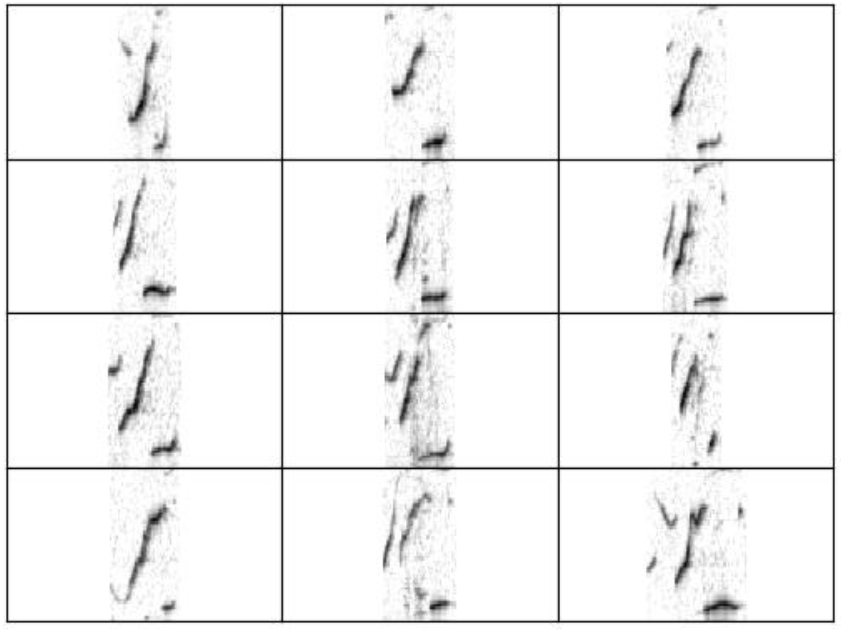
Samples of USVs from the classes ‘c4’ and ‘c5’, USVs with 4 and 5 jumps, respectively. *BootSnap* assigned 68% and 32% of the total members of these two classes to the 2 and 3-jump included USVs, respectively. DSQ assigned the members of the classes ‘c4’ and ‘c5’ mostly to the 2 and 3-jump included USVs and ‘ui’. Although the class ‘ui’ might be relatively similar to the ‘c4’ and ‘c5’ classes based on visual inspection, there is no jump in this class.

